# Coordinate Enhancer Reprogramming by GATA3 and AP1 Promotes Phenotypic Plasticity to Achieve Breast Cancer Endocrine Resistance

**DOI:** 10.1101/767871

**Authors:** Mingjun Bi, Zhao Zhang, Pengya Xue, Karen Hernandez, Hu Wang, Xiaoyong Fu, Carmine De Angelis, Zhen Gao, Jianhua Ruan, Victor X. Jin, Qianben Wang, Elisabetta Marangoni, Tim Hui-Ming Huang, Lizhen Chen, Christopher K. Glass, Wei Li, Rachel Schiff, Zhijie Liu

## Abstract

Acquired therapy resistance is a major problem for anticancer treatment, yet the underlying molecular mechanisms remain unclear. Using an established breast cancer cellular model for endocrine resistance, we show that hormone resistance is associated with enhanced phenotypic plasticity, indicated by a general downregulation of luminal/epithelial differentiation markers and upregulation of basal/mesenchymal invasive markers. Our extensive omics studies, including GRO-seq on enhancer landscapes, demonstrate that the global enhancer gain/loss reprogramming driven by the differential interactions between ERα and other oncogenic transcription factors (TFs), predominantly GATA3 and AP1, profoundly alters breast cancer transcriptional programs. Our functional studies in multiple biological systems including culture and xenograft models of MCF7 and T47D lines support a coordinate role of GATA3 and AP1 in enhancer reprogramming that promotes phenotypic plasticity and endocrine resistance. Collectively, our study implicates that changes in TF-TF and TF-enhancer interactions can lead to genome-wide enhancer reprogramming, resulting in transcriptional dysregulations that promote plasticity and cancer therapy-resistance progression.

## INTRODUCTION

Cancer progression, by which cancer cells adjust themselves to achieve resistance to targeted therapies, is a persistent challenge in cancer treatment. Extensive studies on this phenomenon have revealed that cancer cells can escape targeted therapy through one of the following mechanisms: mutation of the drug targets ^1^; activation of alternate pathways that restore the downstream targets that are inhibited by initial drug blockade ^2^; a recently identified plasticity mechanism by which cell phenotypic plasticity drives therapy resistance ^3–6^.

Breast cancer is a heterogeneous disease. Based on gene expression profiling, breast tumors can be divided into luminal and basal-like subtypes ^7, 8^. The luminal subtype cancers express many luminal epithelial markers including KRT18, GATA3, and PGR. Estrogen (E_2_ or 17β-estradiol) and its nuclear receptor ERα are critical for luminal breast cancer development ^8^. In contrast, the basal-like subtype cancers, which are associated with aggressive pathologic features, are characterized by high expression of mesenchymal genes such as EGFR and many proliferation-related genes ^7^. The luminal subtype cancers consist of 75% of all breast cancers and typically benefit from targeted therapies with drugs that impinge on E_2_/ERα signaling, such as tamoxifen, fulvestrant or aromatase inhibitors ^9, 10^. However, when patients with ERα-positive breast cancer receive endocrine therapies for a period of 5 years, more than 30% of these patients eventually develop resistance and disease recurrence ^9, 10^. As loss of ERα expression during therapies or metastatic progression is only found in ≤10% of patients ^9^, this hormone-receptor pathway remains a major research focus for additional targeted therapies. Substantial evidence suggests that changes of components along the ERα axis, such as mutations or altered expression of ERα itself or ERα-interacting cofactors, may reprogram the ERα-mediated transcriptome that underlie the development of endocrine resistance ^11–13^. Differential ERα binding is also associated with clinical breast cancer progression ^14^. However, the underlying molecular mechanisms of transcriptome transitions mediated by ERα during breast cancer progression and endocrine resistance are not well known.

Enhancers are important distal DNA regulatory elements that control temporal- or spatial-specific gene expression patterns during development and other biological processes ^15–17^. Dysregulation of enhancer function is involved in many diseases, particularly in cancers. Our previous genome-wide ChIP-seq studies have revealed that E_2_/ERα regulates its target gene expression program primarily through binding at distal enhancers to dictate cell growth and endocrine response ^18, 19^. Emerging evidence has implicated the link of epigenetic alterations of ERα-bound enhancers to hormone resistance and cancer invasion ^14, 20^. Thus, investigating oncogenic mechanisms that lead to the alterations of the ERα cistrome is critical for both understanding cancer progression and identifying of potential new cancer therapies.

In this study, we compared matched control and resistant cells lines and identified dramatic changes in ERα-TFs interactions and their cistrome that were associated with genome-wide enhancer gain/loss reprogramming and cancer cell phenotypic plasticity. Among the ERα-interacting TFs, GATA3 and AP1 played a major role in regulating the loss and gain of ERα enhancers respectively. Collectively, our results suggest that altered ERα-TF interactions can reorganize the landscape of ERα-bound enhancers, resulting in gene program transitions that promote plasticity and cancer therapy resistance progression. Our findings provide new insights into the molecular mechanisms underlying plasticity-driven therapy resistance in cancers.

## RESULTS

### Endocrine resistance is associated with plasticity-enhancing transcriptome changes

To study the molecular mechanism of therapy resistance in breast cancer, we used a tamoxifen-resistant (TamR) cell model that was established through long-term culture of ER+ luminal MCF7 parental (MCF7P) cell line in the presence of tamoxifen ^21–23^. We first validated the resistance of TamR cells to 4-OHT and morphological changes. MCF7P cells were sensitive to 4-OHT and displayed a typical epithelial cell-like phenotype and grew in tightly packed cobblestone-like clusters. In contrast, TamR cells were resistant to 4-OHT and began spreading as individual cells, a phenotype similar to mesenchymal cells (Figures S1A and S1B). We next verified that ERα protein levels were comparable between the two lines and that no mutations were detected in ERα gene in TamR cells (Figures S1C and S1D). Thus, tamoxifen resistance in TamR line is not due to altered expression or mutations of ERα.

To evaluate the phenotypic differences at the gene expression level, we performed RNA-seq and identified 1,928 upregulated and 1,899 downregulated genes in TamR when compared to MCF7P (Figure 1A). To test whether the difference in mRNA levels was due to the difference at transcriptional level, we performed Global Run-On coupled with sequencing (GRO-seq) to detect nascent RNAs transcribed by chromatin-associated RNA polymerase ^19^. GRO-seq identified 1,377 upregulated genes and 1,416 downregulated genes in TamR cells (Figure 1A). As shown in the volcano plot, a large percentage of the differentially expressed genes detected by GRO-seq were also captured by RNA-seq (890/1,416 for downregulated genes and 954/1,377 for upregulated genes) (Figure 1A). This result reinforces the notion that transcription regulation plays an important role in the transition of gene expression in tamoxifen resistance, although we cannot exclude the possibility of the post-transcriptional regulation on some genes.

**Figure 1.**
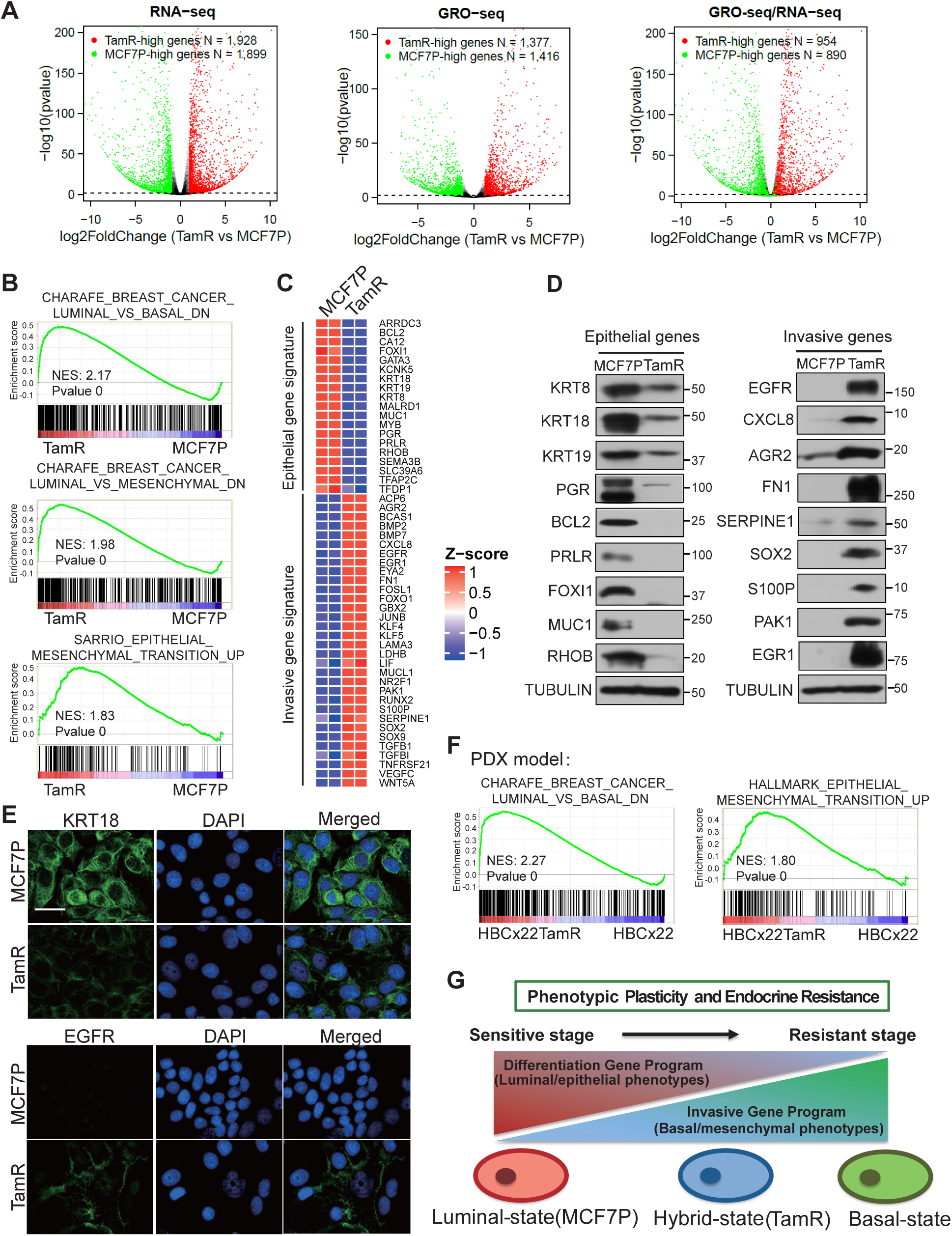
Endocrine resistance is associated with plasticity-enhancing transcriptome changes. (A) Volcano plots showing the genes with differential expression levels in MCF7P and TamR lines detected by RNA-seq (left panel) or GRO-seq (middle panel), and the comparison of their distributions detected by both GRO-seq and RNA-seq (right panel). Each dot represents a gene. In all panels, the green dots are genes significantly downregulated in TamR cells, and the red dots are genes significantly upregulated in TamR cells. In the right panel, the differential genes detected by GRO-seq were re-mapped for their expression changes using RNA-seq data. (B) Gene Set Enrichment Analyses (GSEA) of RNA-seq data for MCF7P and TamR revealing the association of the gene program in TamR cells with the basal and mesenchymal gene signatures. (C) RNA-seq heatmap depiction of selected epithelial marker genes and invasive genes that are differentially expressed in MCF7P and TamR lines. (D) Western blot detection of the protein levels of selected epithelial markers and invasive cancer-associated genes using total cell lysates from MCF7P and TamR lines. (E) Immunofluorescence staining for KRT18 and EGFR in MCF7P and TamR lines. Cell nuclei were stained with DAPI (blue). Scale bar, 30 µm. (F) GSEA analysis of microarray data for paired parental (HBCx22) vs tamoxifen-resistant (HBCx22TamR) PDX tumor samples showing the downregulation of luminal markers and upregulation of EMT markers in tamoxifen-resistant PDX samples. (G) Schematic diagram demonstrating the plasticity-elevating phenotypic transition during the development of endocrine resistance. The luminal breast cancer cells undergo transcriptome transition by reducing differentiation gene program and enhancing invasiveness gene program to achieve resistance.

An unbiased Gene Set Enrichment Analysis (GSEA) ^24^ of our transcriptome data demonstrated that the upregulated genes in TamR cells were significantly enriched in the basal, mesenchymal, and epithelial-to-mesenchymal-transition (EMT) gene signature sets (Figure 1B). These results are consistent with the invasive phenotype observed in TamR cells in previous xenograft studies ^23, 25, 26^. Interestingly, while many basal/mesenchymal invasive genes were upregulated, many of the luminal/epithelial marker genes were downregulated in TamR (Figures 1C, S1E and S1F). We confirmed the downregulation of key luminal/epithelial differentiation genes and the upregulation of invasiveness-associated basal/mesenchymal genes with RT-qPCR (Figures S1G and S1H), Western blotting (Figure 1D) and immunofluorescence staining (Figure 1E). Therefore, TamR cells displayed a gene expression profile featured for EMT and hybrid epithelial/mesenchymal phenotypes. Consistent with TamR phenotype, cancer cells at a hybrid epithelial/mesenchymal state are often more invasive or acquire therapy resistance ^3, 27^.

To extend our findings to clinical patient samples, we used our previously established paired patient-derived xenograft (PDX) models for ER+ luminal tumors (parental vs tamoxifen-resistant) (Cottu et al., 2014) and performed GSEA analyses on the gene expression profiles from the paired PDX samples. Consistent with our findings in the culture models, we observed a downregulation of epithelial markers and an upregulation of EMT signature genes during the acquisition of resistance (Figure 1F). Altogether, these data suggest that cancer cells undergo gene expression transition to enhance phenotypic plasticity (Figure 1G), resulting in a more aggressive EMT-like phenotype during the acquisition of therapy resistance.

### Endocrine resistance is associated with global enhancer reprogramming that drives plasticity-enhancing gene expression

Lineage-specific regulatory elements, especially enhancers, drive cell type- and tissue-specific transcriptional programs ^17^. To evaluate whether the transcriptome changes in endocrine resistance are caused by altered enhancer landscape, we performed H3K4me1 and H3K27ac ChIP-seq analyses of MCF7P and TamR cells. We started with H3K27ac, which marks both active enhancers and promoters, and identified 31,483 and 33,920 H3K27ac peaks in MCF7P and TamR lines respectively. Approximately 50% of these H3K27ac peaks were located at active gene promoters in both cell lines (Figure S2A). The promoters with stronger H3K27ac peaks in TamR than in MCF7P were primarily those of the upregulated genes identified in RNA-seq and GRO-seq. Conversely the promoters with weaker H3K27ac peaks were found at the downregulated genes in TamR cells (Figure S2B). These results further support that transcriptional regulation is the major cause of the altered gene expression in TamR cells.

With ChIP-seq of enhancer markers H3K4me1 and H3K27ac, we identified 7,533 MCF7P-specific enhancers (*LOSS* enhancers) and 10,679 TamR-specific enhancers (*GAIN* enhancers), and 9,896 enhancers shared in both cell lines (*COMMON* enhancers) (Figure S2C). Our ERα ChIP-seq showed that a large portion of these enhancers were ERα-bound enhancers (3,317/7,533 for *LOSS*; 4,450/10,679 for *GAIN*; 6,729/9,896 for *COMMON*) (Figure 2A), suggesting that ERα enhancers are the major contributors for tamoxifen resistance-associated epigenetic changes. Thus, our subsequent studies focused on ERα-bound *LOSS* and *GAIN* enhancers. To validate that these regions are accessible and active enhancers, we profiled chromatin accessibility using ATAC-seq and performed ChIP-seq for the histone acetyltransferase P300. As expected, the *GAIN* ERα-bound enhancers displayed strong P300 binding and high chromatin accessibility in TamR cells, whereas the *LOSS* ERα-bound enhancers have strong P300 binding and high chromatin accessibility only in MCF7P cells (Figure 2A). These results suggest that some of the ERα-bound active enhancers are lost while others are established during the acquisition of tamoxifen resistance.

**Figure 2.**
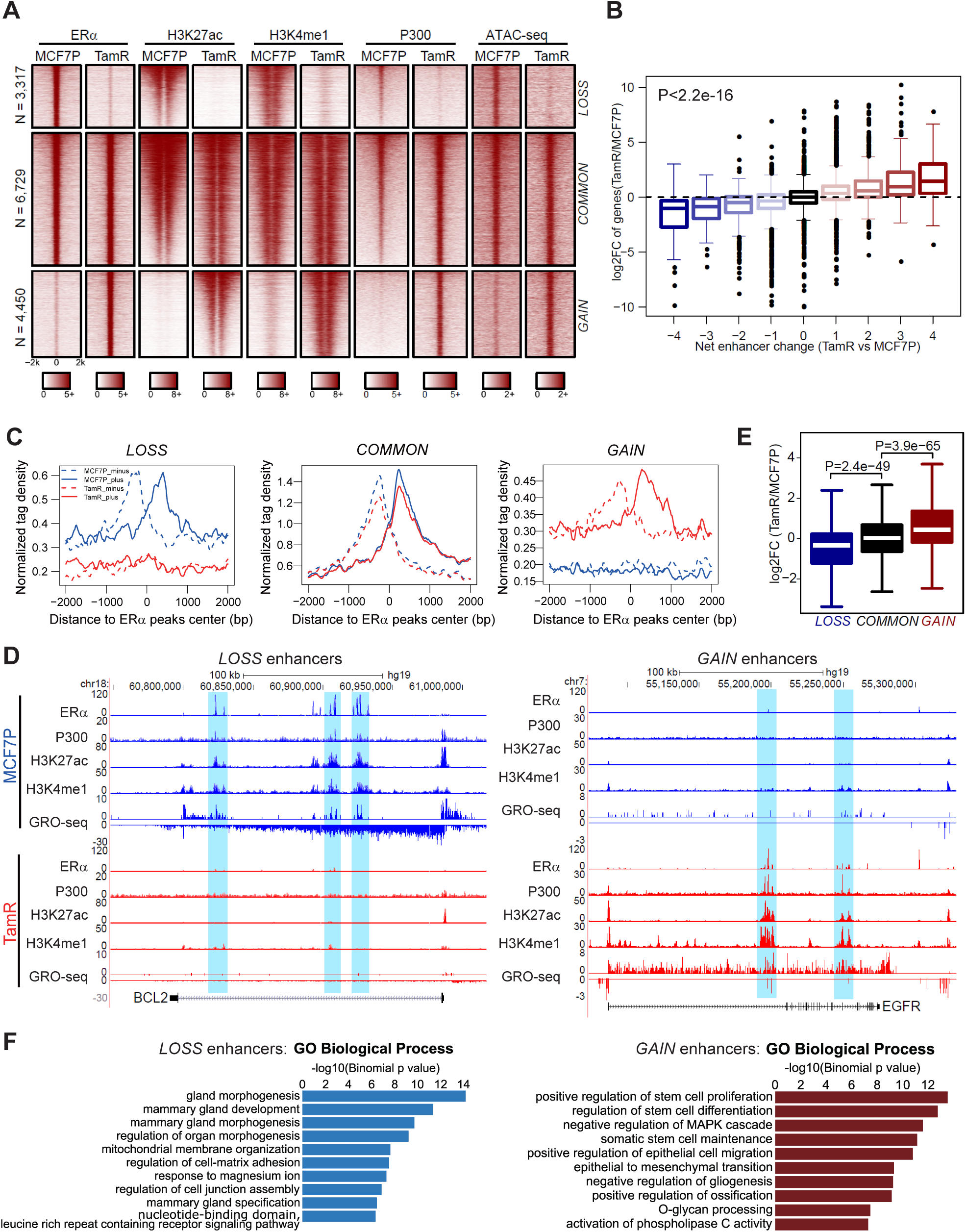
Endocrine resistance is associated with global enhancer reprogramming that drives plasticity-enhancing gene expression. (A) Heatmaps of ERα, H3K4me1, H3K27ac, and P300 ChIP-seq data in both MCF7P and TamR lines demonstrating the difference in enhancer landscape for the three groups of ERα-bound enhancers (*LOSS*, *COMMON* and *GAIN*). Chromatin accessibility profiled by ATAC-seq at the corresponding genomic regions is also shown on the right. (B) Integration of RNA-seq and ChIP-seq data to correlate changes in gene expression with enhancer gain/loss. Box plots showing log_2_(Fold Change) of gene expression for all the genes stratified by the net enhancer change (total number of TamR-specific enhancers minus total number of MCF7P-specific enhancers) within 200 kb from the TSS site of each gene. (C) Aggregate plots of the normalized GRO-seq tag density at *LOSS*, *COMMON* and *GAIN* enhancers in MCF7P (blue) and TamR (red) lines showing the correlation between enhancer activation and the transcription of non-coding enhancer RNA (eRNA). The dashed line represents the minus strand and solid line indicates the plus strand of eRNA. (D) Genome browser views of ChIP-seq and GRO-seq signals at several representative ERα *LOSS* and *GAIN* enhancers and their target genes *BCL2* and *EGFR*. Transcriptional activities (GRO-seq peaks) at enhancers (shaded areas) and their target gene bodies positively correlated with each other, and both associated with enhancer gain/loss. (E) Box plot of the fold changes in expression level of genes adjacent to *LOSS*, *COMMON* and *GAIN* enhancers. P values were calculated with wilcoxon rank sum test. (F) GREAT analyses on the annotations of nearby genes of *LOSS* and *GAIN* enhancers. Top ten enriched annotations are shown.

Next, we examined whether enhancer gain/loss was associated with gene expression changes detected by RNA-seq and GRO-seq. We identified neighboring genes of all the enhancers in both cell lines and stratified these genes into nine groups based on the net enhancer number change within 200 kb of the TSS of each gene: +1 stands for 1 net gained enhancer in TamR; -1 stands for 1 net lost enhancer in TamR; and +0 means no enhancer shift. We observed a high correlation between net enhancer change and gene expression change (Figure 2B), i.e. genes associated with the highest number of net gained enhancers are the most upregulated genes and vice versa. Furthermore, our GRO-seq data demonstrated that the eRNA transcription profiles were consistent with enhancer gain/loss events (Figure 2C), and that the transcriptional activities at enhancers and target gene bodies positively correlated, exemplified by *BCL2* and *EGFR* gene loci (Figure 2D). Annotation analysis also revealed that ERα enhancer gain/loss was associated with the expressional changes of their target genes (Figure 2E). Together, these results suggest that enhancer gain/loss reprogramming accounts for the alterations in gene expression associated with endocrine resistance.

We then used Genomic Regions Enrichment of Annotations Tool (GREAT) ^28^ to interpret functions of genes associated with lost or gained enhancer group. We found that genes associated with *LOSS* enhancer group (MCF7P-specific) were highly enriched for signatures of mammary gland development and morphogenesis, while the genes associated with *GAIN* enhancer group (TamR-specific) were enriched for functions of stem cell proliferation and EMT (Figure 2F). We further used GSEA of our RNA-seq data to interrogate the oncogenic gene signatures from MSigDB database, and identified enriched terms in TamR cells as MEK1, EGFR and ERBB2 pathways (Figure S2D), which are known to regulate endocrine resistance in breast cancer ^29, 30^. Therefore, these data support that enhancer reprogramming triggers gene expression transition and promotes endocrine resistance.

Super-enhancers (SEs) are clusters of enhancers and possess strong enhancer activities and regulate genes with prominent roles in cell identity or diseases ^15^. We have previously demonstrated that many SEs in MCF7 breast cancer cells contain individual ERα-bound enhancers ^19^. To check whether the shifts of ERα-bound enhancers also cause SE reprogramming during endocrine resistance transition, we used H3K27ac signal to rank all enhancers and identified 436 SEs in MCF7P cells and 703 SEs in TamR cells (Figure S2E). Among these SEs, we identified 149 *LOSS* SEs (MCF7P-specific), and 158 *GAIN* SEs (TamR-specific). Notably, alterations of SEs also positively correlated with expressional changes of their nearest target genes (Figure S2F), exemplified by several key luminal/epithelial maker or basal/mesenchymal marker genes (Figure S2G), implicating potential roles of SE reprogramming in gene regulation during cancer progression.

Collectively, our data demonstrate that the acquisition of tamoxifen resistance is associated with global transformation of enhancer chromatin landscapes, rearrangement of ERα occupancy and transition of gene expression.

### Altered interactions between ERα and GATA3/AP1 and their binding on ERα enhancers are associated with enhancer gain/loss reprogramming

We sought to understand the molecular mechanisms underlying enhancer reprogramming during hormone resistance acquisition. We first performed *de novo* motif searches for the three groups of enhancers (*LOSS, COMMON* and *GAIN*). Consistent with its role as a pioneer factor, FOXA1 motif was enriched in all groups (Figure 3A). We also identified the enrichment of the GATA3 and AP2γ motifs on the *LOSS* sites, and the RUNX2 and JUN motifs on the *GAIN* sites (Figure 3A). To examine the distribution pattern of these motifs on all ERα-bound enhancers in both cell lines, we ranked all enhancers based on the ratio of ERα binding strength in the two cell lines (TamR vs MCF7P) and allocated motif occurrence frequency for these enhancers ranked from TamR-high (*GAIN* enhancers) to MCF7P-high (*LOSS* enhancers). The results were consistent with our *de novo* motif analyses above. FOXA1 motif was uniformly distributed across all enhancers (Figure 3B). AP2γ and GATA3 motifs were both enriched in MCF7P-high enhancers, while RUNX2 and AP1 motifs were enriched in TamR-high enhancers (Figure 3B). Notably, ERE motif was highly enriched in the enhancers with similar levels of ERα binding strength in both cell lines (*COMMON* enhancers) (Figure 3B), suggesting that the enhancers undergoing *GAIN*/*LOSS* reprogramming are primarily non-ERE ERα-bound enhancers. Since ERα can bind to the chromatin either in *cis* (directly binds to ERE motif) or in *trans* (binds to chromatin through tethering to other TFs) ^31, 32^, ERα might bind to these *GAIN*/*LOSS* enhancers in *trans* through protein-protein interaction.

**Figure 3.**
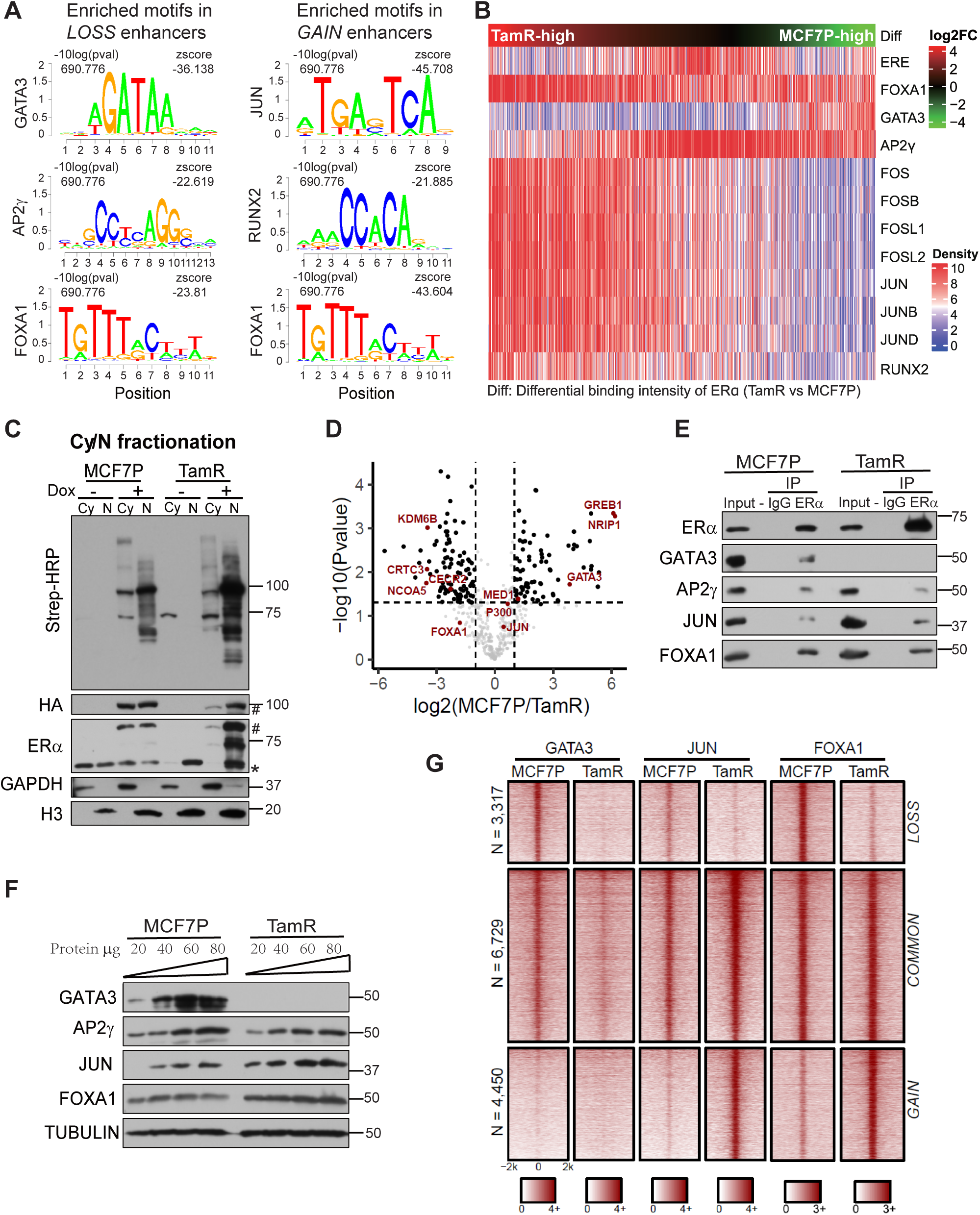
Altered interactions between ERα and GATA3/AP1 and their binding on ERα enhancers are associated with enhancer gain/loss reprogramming. (A) Enriched motifs in different enhancer groups. (B) Heatmap of motif densities for the listed TFs at all *LOSS*, *COMMON* and *GAIN* enhancers that are arranged according to the differential binding intensities of ERα measured by the ratio of normalized ERα reads in TamR relative to MCF7P. A motif is considered occurred in an enhancer if the p-value for the region with maximum score is less than 1e-4 by FIMO scanning of this enhancer. (C) Western blots confirming the inducible expression and *in vivo* biotinylation in the established ERα-BioID tet-on stable cell lines. The fractionation of cytoplasmic (Cy) and nuclear (N) fractions of MCF7P or TamR cells was confirmed with Western blots for GAPDH (cytoplasm-specific marker) and Histone H3 (nucleus-specific marker). The doxycycline-induced ERα-BirA*-HA fusion protein expression was detected by antibodies recognizing HA, ERα (the endogenous ERα was labeled with * and the tagged exogenous ERα was labeled with #). Proteins biotinylated by ERα-BirA* were detected using streptavidin-HRP blot. (D) Volcano plot showing the log2(LFQ) value for ERα-associated proteins identified in all four BioID replicates. Several ERα-interacting proteins are highlighted in red. (E) GATA3, AP2γ, FOXA1 and JUN bind to endogenous ERα in MCF7P and TamR cells. Endogenous ERα was immunoprecipitated using anti-ERα antibody, and its interactions with different TFs were confirmed with immunoblots. IgG was used as a negative control. (F) Western blot analyses of the protein levels of GATA3, AP2γ, JUN and FOXA1 in MCF7P and TamR cells. Tubulin was used as a loading control for different samples. (G) Heatmaps of GATA3, JUN and FOXA1 ChIP-seq data in both MCF7P and TamR lines demonstrating their occupancy on *LOSS*, *COMMON* and *GAIN* enhancers.

We considered whether ERα interacted with different TFs in different contexts to facilitate enhancer *GAIN*/*LOSS* reprogramming. We used a powerful *in vivo* proximity proteomics approach BioID ^33^ to identify context-specific ERα cofactors. We generated MCF7P and TamR tet-on stable lines expressing HA-tagged ERα fused with a mutant biotin ligase (BirA*), which promiscuously biotinylates interacting/neighboring proteins *in vivo* (Figure S3A). To identify ERα cofactors (i.e. proteins biotinylated by the BirA*-ERα), streptavidin beads used for BioID pull-down from nuclear lysate were digested and subjected to liquid chromatography-tandem mass spectrometry (LC-MS/MS) (Figure 3C). MS identified 475 ERα-associated proteins in MCF7P and TamR cells with at least two unique peptides. The robustness of our BioID experiments was demonstrated by the recovery of many previously known ERα-interacting partners including TFs such as FOXA1, AP2γ, GATA3, and epigenetic co-regulators such as P300, MED1, NCOA3, GREB1 and NRIP1 (Figure 3D). Consistent with our hypothesis, we identified context-specific ERα-cofactor interactions: its interactions with GREB1, NRIP1, and GATA3 were only detected in MCF7P cells, and its interaction with NCOA5 and FOXA1 were stronger in TamR cells (Figure 3D). Indeed, loss of interaction between ERα and GREB1 ^34^ and overexpression of FOXA1 ^21^ are known to associate with hormone resistance, further supporting the authenticity of our BioID data. Several AP1 family TFs were among the identified ERα-interacting proteins (Figure 3D), we later on chose JUN to study AP1 function on *GAIN* enhancers, as it is a common component of JUN/FOS and JUN/JUN dimers ^35^.

We next confirmed the context-specific interaction between ERα and some of the BioID-identified cofactors with co-immunoprecipitation (co-IP) experiments using ERα antibody. Based on the IP results, ERα showed weaker interaction strength with GATA3 and AP2γ in TamR, while its interactions with FOXA1 and JUN were stronger (Figure 3E). This could be partly due to differential expression of the cofactors, as we detected lower level of GATA3 and AP2γ, but higher level of FOXA1 and JUN in TamR (Figure 3F). Phosphorylation of serines 63 and 73 in the transactivation domain of JUN is known to enhance its transcriptional activity ^36^. We also detected higher level of phosphorylated JUN in TamR (Figure S3B), indicating that tamoxifen resistance is associated with both higher expression and activity of JUN. These results suggest that in different contexts ERα interacts with different TF partners, which may promote enhancer reprogramming by facilitating the recruitment of ERα to different sets of enhancers.

Thus, we decided to test whether these ERα-interacting TFs could bind to chromatin and regulate enhancer gain/loss reprogramming. We performed ChIP-seqs for GATA3, JUN and FOXA1 in MCF7P and TamR cell lines and revealed context-dependent binding patterns for GATA3 and JUN. While FOXA1 bound to all three groups of enhancers and *COMMON* enhancers recruited all three TFs, the 3,317 MCF7P-specific *LOSS* enhancers recruited GATA3 and excluded JUN, and conversely, the 4,450 TamR-specific *GAIN* enhancers recruited JUN and excluded GATA3 (Figures 3G and S3C-S3E), indicating the association of GATA3 with *LOSS* enhancers and JUN with *GAIN* enhancers. Together, these results suggest that reduced GATA3 expression in TamR may lead to loss of GATA3-bound enhancers, while increased expression and activity of JUN in TamR could be involved in *de novo* establishment of JUN-bound enhancers. These results support a model that FOXA1 is a pioneer factor for all enhancers and GATA3 and AP1 are context-specific regulators working together with FOXA1 for enhancer reprogramming during the acquisition of tamoxifen resistance.

### GATA3 is required for maintenance of *LOSS* enhancers and expression of epithelial makers

GATA3 is a key determinant of luminal-type breast cancers, and can inhibit metastases by inducing epithelial fate and suppressing mesenchymal fate simultaneously ^37, 38^. As GATA3 expression was greatly reduced in TamR cells (Figure 3F), we asked whether GATA3 expression was affected by DNA methylation. Using published genome-wide methylation profiles derived from three different types of endocrine-resistant MCF7 cells ^39^, we identified a CpG-containing area at 5’ end of GATA3 gene locus with a remarkably higher methylation level in all three different resistant lines than in the parental cells (Figure S4A). Similarly, our pyrosequencing revealed significantly higher level of DNA methylation in the same area in our TamR cells than MCF7P cells (Figure S4A), suggesting that GATA3 gene hypermethylation might be a general phenomenon associated with endocrine resistance. We also found that treatment of DNA methyltransferase inhibitor 5-Aza-2’-deoxycytidine (5-Aza) was able to enhance *GATA3* mRNA expression in TamR but not MCF7P cells (Figure S4B), confirming the effect of DNA hypermethylation on GATA3 silencing in TamR cells. Furthermore, TCGA methylome data for breast cancers showed that DNA methylation signals at this particular locus negatively correlated with GATA3 expression and positively correlated with invasiveness among different tumor subtypes (Figure S4C). Altogether, these data suggest that DNA methylation-mediated GATA3 silencing might promote tamoxifen resistance and cancer invasive progression.

So far, our multiple lines of evidence from BioID proteomics, ChIP-seq, and GATA3 expression all point to the key role of GATA3 in maintaining the MCF7P-specific ERα-bound enhancers that are lost in TamR cells. To functionally test this, we generated GATA3-deficient MCF7P cells with small hairpin RNA (shRNA)-mediated knockdown (KD). We found that GATA3 KD dramatically decreased ERα binding signal at these *LOSS* enhancers (Figure 4A). In agreement with a previously proposed role of GATA3 in shaping chromatin architecture in a subset of ERα-bound enhancers ^40^, our data showed that GATA3 depletion led to a great reduction in H3K27ac and eRNA transcription on these *LOSS* enhancers (Figures 4A and S4D), exemplified by *BCL2* and *KCNK5* gene loci (Figure 4B). Conversely, GATA3 overexpression (OE) in TamR cells was able to promote *LOSS* enhancer activation, indicated by the increased H3K27ac level (Figure S4E). Thus, these findings suggest that GATA3 regulates enhancer landscape to maintain ERα binding at these *LOSS* enhancers in MCF7P cells and that epigenetic silencing of GATA3 in TamR cells causes the inactivation of these enhancers.

**Figure 4.**
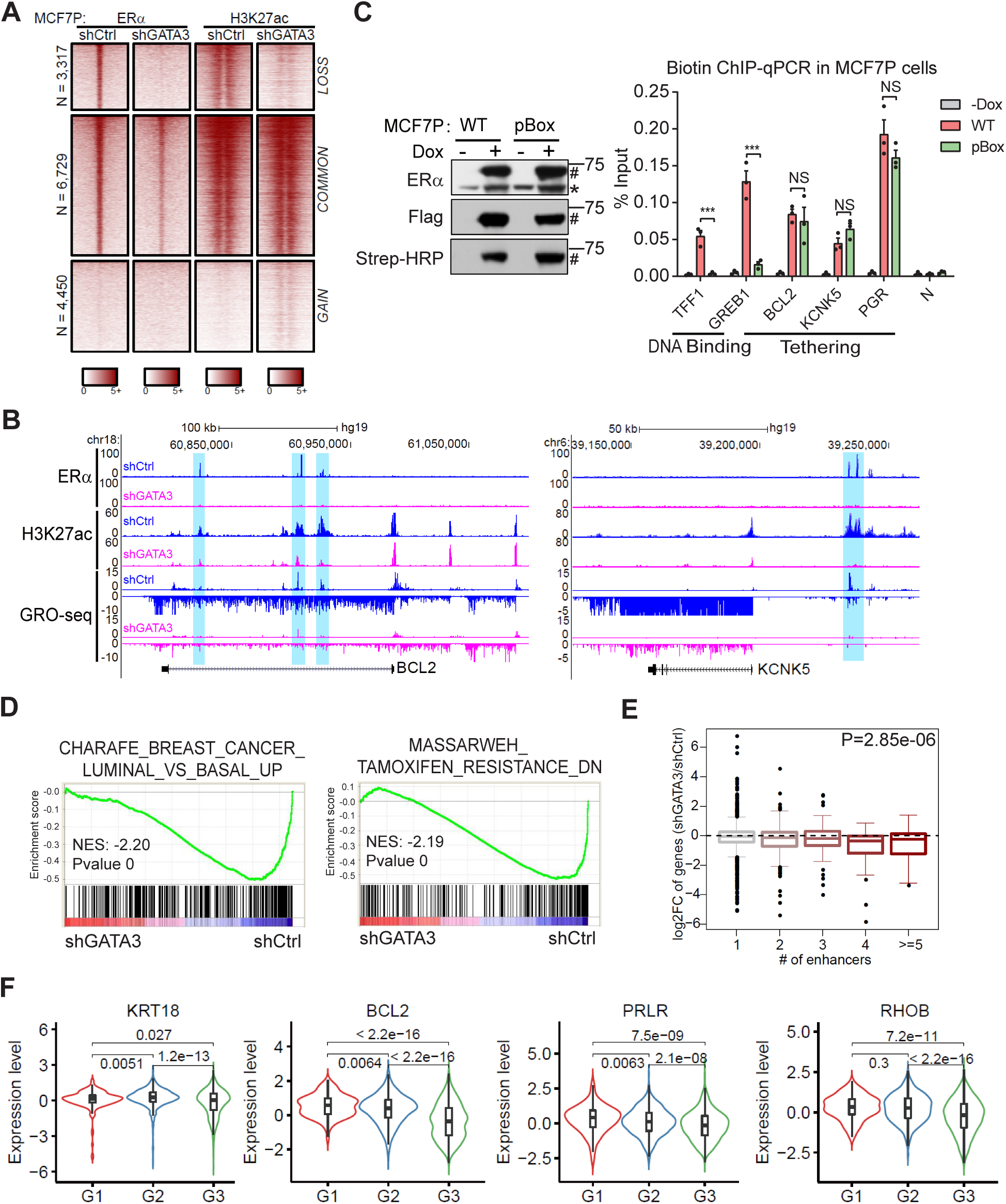
GATA3 is required for maintenance of *LOSS* enhancers and expression of epithelial makers. (A) Heatmaps of ERα and H3K27ac ChIP-seq data at *LOSS*, *COMMON* and *GAIN* enhancers demonstrating that knockdown of GATA3 in MCF7P cells results in enhancer inactivation for the *LOSS* group. (B) Genome browser views of GRO-seq data and ChIP-seq data for ERα and H3K27ac at the *BCL2* and *KCNK5* gene loci demonstrating the depletion of GATA3 in MCF7P cells causes the enhancer inactivation and downregulation of gene expression. (C) Western blots (left) showing doxycycline-induction and *in vivo* biotinylation of BLRP-tagged ERα WT and pBox mutant in MCF7P cells. * and # indicate endogenous and tagged exogenous ERα respectively. ChIP-qPCR showing that the binding of ERα on *LOSS* enhancers is independent of pBox-mediated DNA binding activity, in contrast to its binding on the classical ERE-containing enhancers at *TFF1* and *GREB1* loci (*cis* binding that requires DNA binding ability to bind DNA motif). Data are presented as mean ± SEM. NS, not significant. *** P < 0.001. (D) GSEA analyses on RNA-seq data for MCF7P cells transduced with shCtrl or shGATA3 lentiviruses showing that the signature genes of luminal and tamoxifen-sensitive cancers are downregulated after GATA3 knockdown in MCF7P cells. (E) Integration of RNA-seq and ChIP-seq data to correlate gene regulation effects by GATA3 knockdown and GATA3-bound *LOSS* enhancers in MCF7P cells. Box plots showing GATA3 knockdown effects on these genes stratified by the numbers of nearest *LOSS* enhancers within 200 kb from the TSS site of each gene. Genes associated with higher number of *LOSS* enhancers display more significant downregulation upon GATA3 knockdown in MCF7P cells. (F) Negative correlation between the average gene expression levels of several validated GATA3 direct targets (KRT18, BCL2, PRLR and RHOB) and tumor grades (G1, G2 and G3). TCGA RNA-seq data from breast cancer samples were used in the analyses.

As motif analyses on the *LOSS* enhancers identified low enrichment of ERE motif but high enrichment of GATA3 motif, we reasoned that ERα might be recruited to these GATA3-bound enhancers *in trans* through protein-protein tethering. To test this possibility, we compared wild-type ERα and DNA-binding defective ERα for their occupancy on enhancers. We generated MCF7P stable lines expressing BirA that can recognize and biotinylate BLRP-tagged proteins *in vivo* ^19^, These cell lines were induced to express BLRP-tagged wild-type ERα or mutant ERα with disrupted pBox domain for *in vivo* biotinylation and streptavidin pulldown (Figure 4C). Our ChIP-qPCR data showed that DNA-binding deficient ERα lost its binding to the ERE-containing classic *COMMON* enhancers located at *TFF1* and *GREB1* gene loci, but maintained its binding to the GATA3 motif-enriched *LOSS* enhancers associated with luminal/epithelial marker genes *BCL2*, *KCNK5*, and *PGR* (Figure 4C). Therefore, these results suggest that the recruitment of ERα to *LOSS* enhancers might be via tethering to GATA3 or other TFs such as AP2γ, which can directly modulate the landscape of the MCF7P-specific ERα enhancers.

To evaluate the impact of GATA3-mediated enhancer reprogramming on gene regulation, we performed RNA-seq in MCF7P cells treated with GATA3 shRNA and identified 255 downregulated genes and 341 upregulated genes. Interestingly, we found that the most dramatically downregulated genes are highly enriched for luminal/epithelial gene signature and tamoxifen-sensitive gene signature (Figures 4D and S4F). This result is consistent with the GREAT analyses of *LOSS* enhancers above (Figure 2F). Furthermore, the correlation between the number of *LOSS* enhancers within 200 kb from the TSS site of a gene and the degree of gene expression downregulation by GATA3 KD implies that GATA3 is required for activation of a subset of luminal/epithelial genes through regulating *LOSS* enhancers (Figure 4E). We also confirmed that the protein levels of several key luminal/epithelial genes (*KRT18*, *BCL2*, *PRLR*, *FOXI1* and *RHOB*) were significantly decreased upon GATA3 KD (Figure S4G). More importantly, overexpressing GATA3 in TamR cells was sufficient to re-sensitize the cells to tamoxifen treatment (Figures S4H). Given that lower levels of GATA3 in human breast cancer are associated with more aggressive tumors and worse prognosis ^41^, we considered whether the target genes of GATA3-controlled *LOSS* enhancers can serve as biomarkers of tumor invasiveness and have prognostic values. To test this idea, we analyzed TCGA data from annotated and transcriptionally profiled breast cancer samples. Indeed, we observed a negative correlation between the expression levels of GATA3 direct targets and tumor grades (Figure 4F). Further supporting this notion, the expression level of BCL2, a direct target of GATA3, is found to positively correlate with relapse free survival (RFS) in the breast cancer patients receiving endocrine therapy (Figure S4I). In summary, our data suggest that epigenetic silencing of GATA3 results in specific loss of ERα-bound enhancers and downregulation of a subset of luminal/epithelial genes to promote endocrine resistance and invasive phenotypes.

### AP1-mediated *GAIN* enhancer activation promotes hormone resistance-associated gene program and phenotypes

Having shown the association of AP1 with the *GAIN* enhancers (Figure 3), we next investigated AP1 function in *GAIN* enhancer activation. AP1 is known to respond to tumorization-associated signals and regulate proliferation, survival, and invasiveness of tumor cells in various cancers ^35, 42^. Breast cancers with acquired tamoxifen resistance displayed enhanced transcriptional activity of AP1 ^43, 44^, but the underlying mechanism is not clear. A previous work has shown that AP1 potentiates chromatin accessibility and facilitates the binding of glucocorticoid receptor to chromatin ^45^. Thus, we evaluated whether *GAIN* enhancers that are silent in MCF7P cells could be activated by JUN OE. We found that Dox-induced JUN OE significantly elevated ERα occupancy and H3K27ac level at *GAIN* regions in MCF7P cells (Figure 5A). It also upregulated a set of genes with TamR-associated basal/mesenchymal gene signatures (Figures 5B and S5A). The upregulation of well-known endocrine resistance-related genes such as *EGFR*, *CXCL8*, *AGR2*, *S100P*, and *PAK1*, was confirmed at protein levels by immunoblots (Figure S5B). Furthermore, we observed a strong correlation between the number of *GAIN* enhancers within 200 kb from the TSS site of a gene and the degree of gene expression upregulation by JUN OE (Figure 5C). These results support that JUN is a key driver of *GAIN* enhancer reprogramming and invasive phenotype-associated gene expression during endocrine resistance progression.

**Figure 5.**
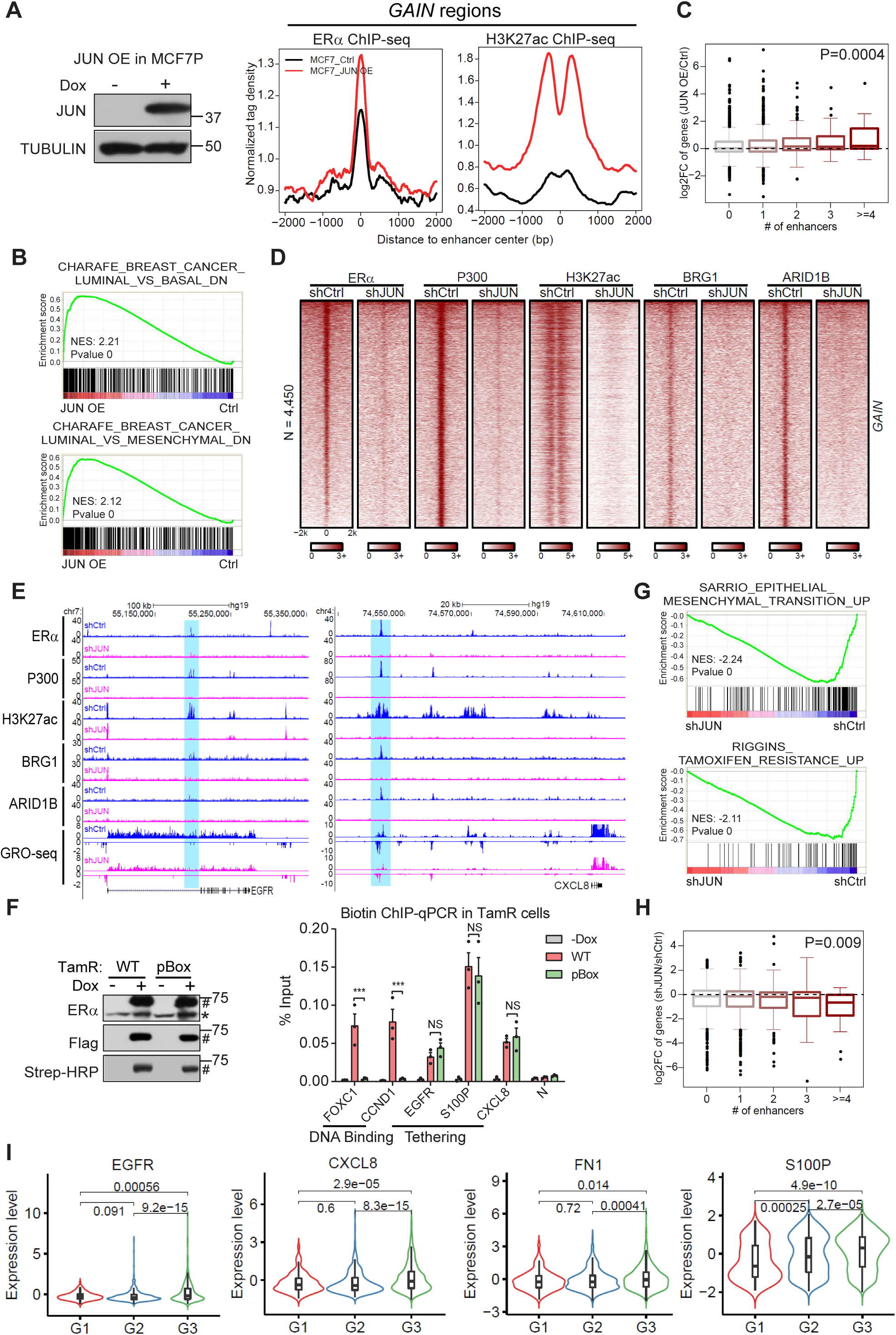
AP1-mediated *GAIN* enhancer activation promotes hormone resistance-associated gene program and phenotypes. (A) Aggregate plots showing the normalized tag density of ERα and H3K27ac ChIP-seq data at *GAIN* enhancers (right). The results suggest overexpression of JUN in MCF7P cells can activate these silent *GAIN* enhancers. Western blot confirms the doxycycline-induced expression of JUN in MCF7P cells (left). Tubulin was used as a loading control. (B) GSEA analyses of RNA-seq data for MCF7P cells with or without JUN OE showing that basal and mesenchymal gene signatures were upregulated upon JUN OE in MCF7P cells. (C) Integration of RNA-seq and ChIP-seq data to correlate gene regulation effects by JUN overexpression (OE) and JUN-bound *GAIN* enhancers in MCF7P cells. Box plots representation of JUN OE effects on expression changes of genes stratified by the numbers of nearest JUN-bound *GAIN* enhancers within 200 kb from the TSS site of each gene. Genes associated with higher number of JUN-bound *GAIN* enhancers show higher levels of upregulation upon JUN OE in MCF7P cells. (D) Heatmaps of ERα, P300, H3K27ac, BRG1 and ARID1B ChIP-seq data at *GAIN* enhancers in TamR cells demonstrating that JUN KD greatly deactivates *GAIN* enhancers and causes the loss of chromatin remodeling factors including BRG1 and ARID1B from these enhancers. (E) Genome browser views of GRO-seq data and ChIP-seq data for ERα, P300, H3K27ac, BRG1 and ARID1B at the *EGFR* and *CXCL8* gene loci. Depletion of JUN in TamR cells leads to enhancer inactivation and transcriptional downregulation. (F) Western blots (left) showing that the doxycycline-induction and *in vivo* biotinylation of BLRP-tagged ERα (WT and pBox mutant) in TamR cells. * and # indicate endogenous and tagged exogenous ERα respectively. Biotin ChIP-qPCR shows that ERα binding on *GAIN* enhancers is via *trans* binding and is independent of its DNA binding ability. In contrast, its binding to the classical ERE-containing enhancers at *FOXC1* and *CCND1* loci is via *cis* binding and requires DNA binding ability to bind DNA motif. Data are presented as mean ± SEM. NS, not significant. *** P < 0.001. (G) GSEA analyses of RNA-seq data showing that EMT and tamoxifen resistance related gene signatures were downregulated upon JUN KD in TamR cells. (H) Integration of RNA-seq and ChIP-seq data to investigate gene regulation effects by JUN KD on JUN-bound *GAIN* enhancers in TamR cells. Box plots showing JUN KD effects on genes stratified by the numbers of nearest JUN-bound *GAIN* enhancers within 200 kb from the TSS site of each gene. Genes associated with higher number of JUN-bound *GAIN* enhancers display more significant downregulation upon JUN KD in TamR cells. (I) The average gene expression values of several validated JUN direct targets (*EGFR*, *CXCL8*, *FN1* and *S100P*) positively correlate with tumor grades (G1, G2 and G3). TCGA RNA-seq data from breast cancer samples were used in the analyses.

To further test the critical role of JUN in regulating *GAIN* enhancers, we examined the effect of JUN KD on enhancer chromatin landscape in TamR cells. Remarkably, ERα occupancy at the *GAIN* enhancers was completely diminished upon JUN KD, which paralleled a dramatic reduction in the recruitment of P300 and H3K27ac to the *GAIN* regions (Figure 5D). We also performed ChIP-seq for the components of the BAF chromatin-remodeling complex and observed attenuated occupancy of BRG1 and ARID1B on the *GAIN* sites (Figure 5D). This is consistent with a recent report that AP1 can mediate signal-dependent enhancer selection by recruiting the BAF complex to establish accessible chromatin ^46^. The effects of JUN KD on *GAIN* enhancer landscape were exemplified at the *EGFR* or *CXCL8* gene loci (Figure 5E). Moreover, GRO-seq data revealed a reduction of eRNA transcription from the *GAIN* sites upon JUN KD (Figure S5C). Similar to the enrichment of GATA3 motif on *LOSS* enhancers, AP1 motif was highly enriched on *GAIN* enhancers, with a more significant enrichment than ERE motif (Figure 3B). Using the same BirA-BLRP biotin-tagging approach described above, we showed that DNA-binding activity of ERα was not required for its binding to *GAIN* enhancers ((Figure 5F). Therefore, the recruitment of ERα to *GAIN* enhancers might be via tethering to AP1.

To determine the role of AP1-mediated enhancer reprogramming on gene expression, we performed RNA-seq in TamR cells treated with shJUN and identified 328 upregulated and 1,176 downregulated genes, which included those EMT- and tamoxifen resistance-associated genes (Figures 5G and S5D). Integration of the differential gene expression upon JUN KD and *GAIN* enhancer numbers within 200 kb from the TSS sites of those genes revealed a correlation between the degree of downregulation and the number of their neighboring *GAIN* enhancers (Figure 5H). Downregulation of key cancer invasiveness marker genes, including EGFR, CXCL8, AGR2, FN1, S100P and PAK1, was confirmed at protein level (Figure S5E). Since JUN KD led to downregulation of invasiveness-associated genes, and blockade of AP1 was reported to overcome endocrine resistance in human breast cancer ^47^, we wondered whether JUN depletion was sufficient to affect cancer cell behavior. Indeed, knocking down JUN in TamR cells was able to re-sensitize cells to tamoxifen treatment (Figure S5F). Furthermore, TCGA data from annotated and transcriptionally profiled breast cancer samples showed that the expression levels of several direct target genes of JUN positively correlated with tumor grades (Figure 5I). In addition, we found that higher expression of some of the JUN direct targets, such as *FN1* and *S100P*, is associated with worse relapse free survival (RFS) in the breast cancer patients receiving endocrine therapy (Figure S5G). Taken together, these findings suggest that JUN-mediated enhancer activation promotes resistance-associated gene program and phenotypes.

### Coordinate role of GATA3 and AP1 in promoting TamR-associated enhancer reprogramming and gene expression

Having demonstrated the individual function of GATA3 and JUN in controlling *LOSS* and *GAIN* enhancer activation and gene expression, we were then interested in testing the combined effect of manipulating GATA3 and JUN simultaneously. As GATA3 level is high and JUN expression is low in MCF7P cells, we treated MCF7P cells with either shGATA3 or JUN OE vector, or both. We then performed RT-qPCRs and western blots to determine the expressional changes of epithelial markers (KRT18, BCL2, and PRLR) and invasive markers (EGFR, S100P, and FN1). Compared to single manipulations, double manipulation demonstrated a combined effect in inhibiting epithelial marker expression or in elevating invasiveness-associated genes (Figures S6A and S6B), which was also supported by RNA-seq data under different manipulation conditions in MCF7P cells (Figure 6A). Similar combined effect of GATA3 KD and JUN OE on gene expression was also observed in T47D ER+ luminal breast cancer cells (Figures S6C and S6D). When we examined the H3K27ac profiles in MCF7P cells, we found that GATA3 KD and JUN OE together displayed synergistic effect on both *LOSS* and *GAIN* enhancers, resulting in re-organization of enhancer landscape that mimicked the transition from MCF7P to TamR (Figure 6B). These findings demonstrate that manipulating GATA3 and AP1 simultaneously can more efficiently reprogram enhancer landscape and promote TamR-like properties than manipulating either gene individually. Like in MCF7P cells, simultaneous manipulation of GATA3 and JUN in T47D cells resulted in more profound enhancer LOSS/GAIN reprogramming compared to single factor manipulation (Figure S6E), suggesting that the cooperation between GATA3/AP1 and ERα on enhancers might be a common mechanism to reprogram enhancers in different ER+ breast cancer cell lines.

**Figure 6.**
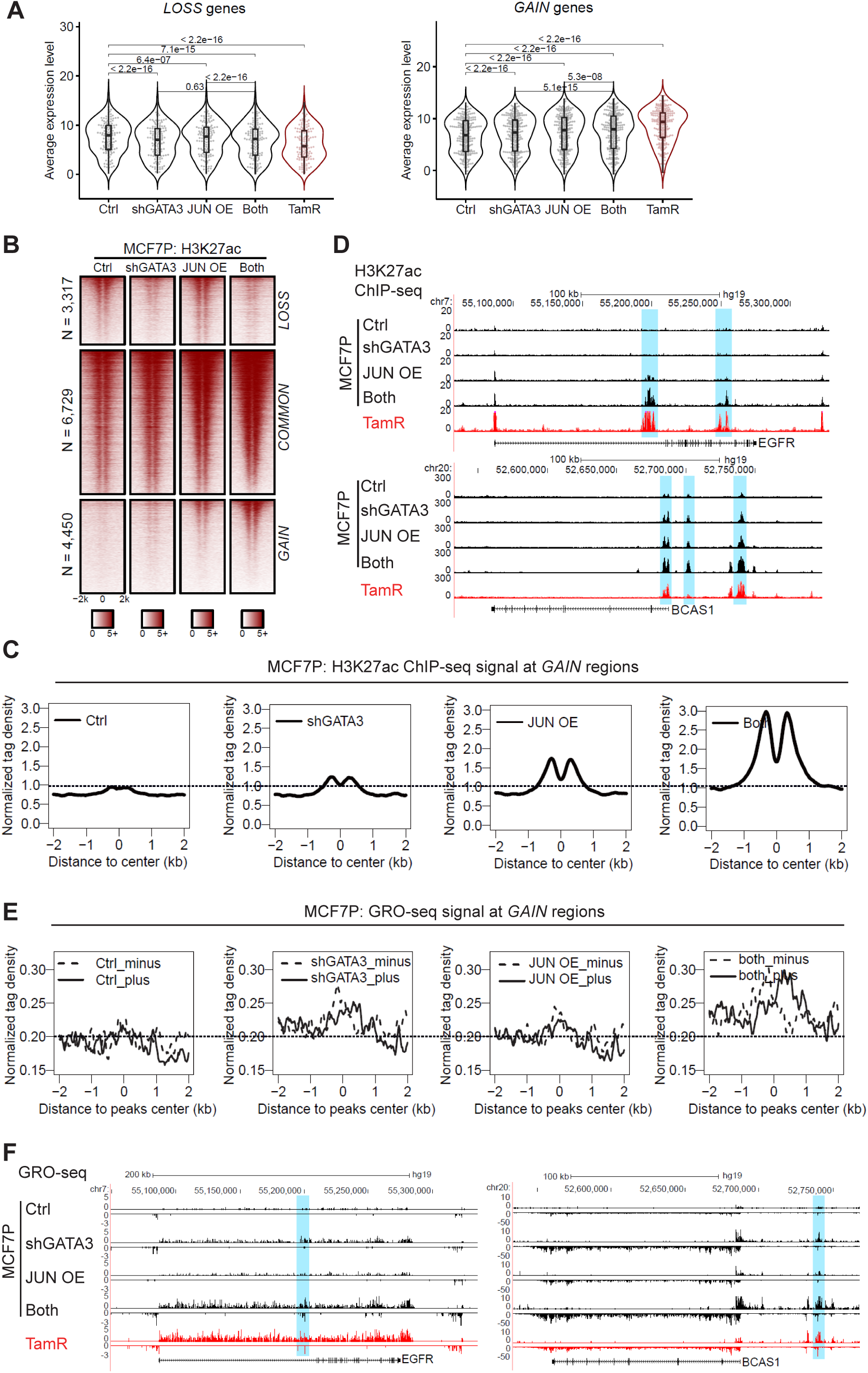
Coordinate role of GATA3 and AP1 in promoting TamR-associated enhancer reprogramming and gene expression. (A) Box plots representation of gene expression in MCF7P cells. Simultaneously depleting GATA3 and overexpressing JUN (“both”) shows a more dramatic effect on the lost and gained gene expression in MCF7P cells compared to manipulating individual gene alone. P values were calculated by Wilcoxon signed rank test. (B) Heatmaps of H3K27ac ChIP-seq data at *LOSS*, *COMMON* and *GAIN* enhancers in MCF7P cells with the indicated treatments. (C) The aggregate plots of the normalized tag densities of H3K27ac ChIP-seq data at *GAIN* enhancers in MCF7P cells with indicated treatments. GATA3 KD and JUN OE demonstrate a synergistic effect on activating *GAIN* enhancers. (D) Genome browser snapshots of H3K27ac ChIP-seq signals at the *EGFR* and *BCAS1* gene loci. GATA3 KD and JUN OE show a synergistic effect. The combined treatment in MCF7P cells creates an enhancer landscape similar to that in TamR cells. (E) The aggregate plots of the normalized GRO-seq tag density at *GAIN* enhancers in MCF7P cells under indicated treatments. GATA3 KD and JUN OE demonstrate a synergistic effect on eRNA transcription. The dashed line represents the minus strand and solid line indicates the plus strand of eRNA. (F) Genome browser snapshots of GRO-seq signals at the *EGFR* and *BCAS1* gene loci in MCF7P cells. MCF7P cells with simultaneous GATA3 KD and JUN OE (“both”) display similar levels of enhancer and gene body activation to TamR cells.

Although GATA3 predominantly regulates *LOSS* enhancer group, we also found its silencing in luminal MCF7P cells led to a slight but significant increase of H3K27ac level on the *GAIN* enhancers (Figure 4A). Inversely, GATA3 OE in TamR cells caused a significant reduction of H3K27ac signal at *GAIN* enhancers (Figure S4E). But we did not observe an effect of JUN OE on *LOSS* enhancers (Figure 6B). In MCF7P cells with JUN OE, GATA3 KD could magnify the AP1-mediated enhancer activation effect on *GAIN* enhancers (Figures 6B and 6C), exemplified by several *GAIN* enhancers at the loci of *EGFR* and *BCAS1* genes (Figure 6D). The effect of GATA3 KD on *GAIN* enhancers was also observed in T47D cells (Figures S6F and S6G). Furthermore, GRO-seq analysis showed that GATA3 KD synergized with JUN OE to promote eRNA transcription from *GAIN* sites (Figures 6E and 6F). These data suggest that during acquisition of endocrine resistance, loss of GATA3 not only leads to inactivation of *LOSS* enhancers, but also results in elevated AP1-mediated activation of *GAIN* enhancers. Since GATA3 does not bind to *GAIN* enhancers (Figure 3G), it might suppress *GAIN* enhancer activation in MCF7P cells via an indirect mechanism.

### Combined effect of GATA3 and AP1 in promoting endocrine resistance and tumor growth *in vitro* and *in vivo*

Given the coordinate role of GATA3 and AP1 in controlling enhancer landscape and gene expression, we next evaluated whether GATA3 and AP1 cooperated to drive endocrine resistance and cancer growth. Compared to either individual manipulation, simultaneous GATA3 KD and JUN OE in MCF7P led to more obvious morphological changes, including cellular elongation with a mesenchymal-like appearance and a dispersed growth pattern (Figure S7A). Similarly, combined effects of GATA3 KD and JUN OE on cell morphology were also observed in T47D cells (Figure S7B). We next examined how simultaneously knocking down GATA3 and overexpressing JUN in MCF7P cells affect their response to tamoxifen. Either GATA3 KD or JUN OE in MCF7P cells was able induce resistance to tamoxifen. Notably, MCF7P cells with simultaneous GATA3 KD and JUN OE displayed a stronger resistant phenotype than cells with individual gene manipulation (Figure 7A). Similar results were obtained in T47D cells (Figure 7B).

**Figure 7.**
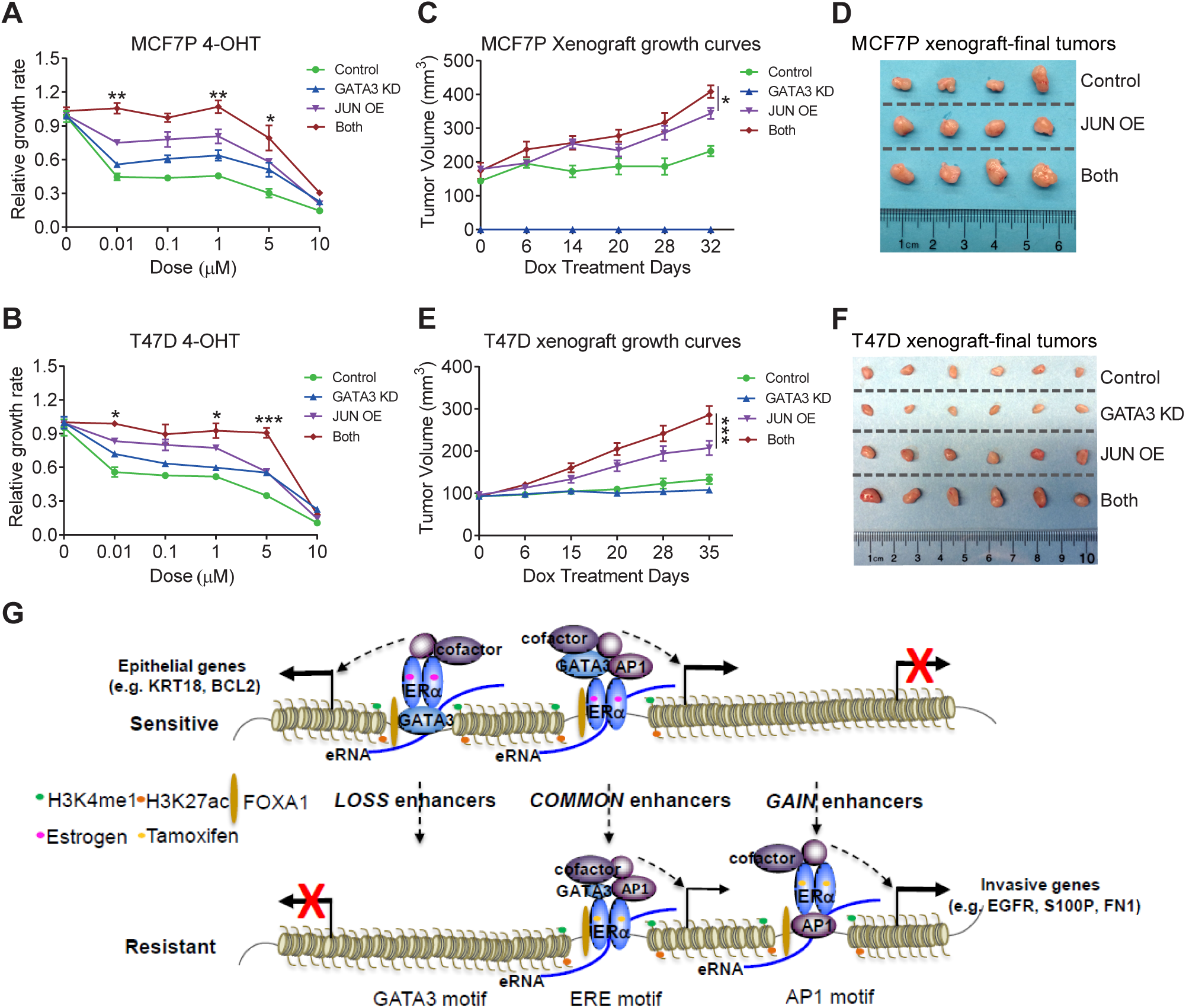
Combined effect of GATA3 and AP1 in promoting endocrine resistance and tumor growth *in vitro* and *in vivo*. (A) Knocking down GATA3 and/or overexpressing JUN in MCF7P cells increases their resistance to 4-OHT. MCF7P stable cell lines expressing shGATA3 and/or JUN were used in the CCK8 assays to measure the relative cell viability after indicated treatments for 5 days. Data are presented as mean ± SEM. *P < 0.05, **P < 0.01 by 2-tailed t test. (B) Knocking down GATA3 and/or overexpressing JUN in T47D cells promotes their resistance to 4-OHT. T47D stable cell lines expressing shGATA3 and/or JUN were used in the CCK8 assays to measure the relative cell viability after indicated treatments for 5 days. Data are presented as mean ± SEM. *P < 0.05, *** P < 0.001 by 2-tailed t test. (C and E) Tumor growth curves of orthotopic xenografts of manipulated MCF7P cells or T47D cells in nude mice (n = 4 per group). MCF7P cells or T47D cells with JUN OE showed enhanced tumor growth, which was further enhanced by GATA3 KD. Tamoxifen subcutaneously injections were performed right after the graft (1 mg/mouse, three times/week). Tumor sizes were measured once a week upon starting doxycycline (administrated in water). Data are presented as mean ± SEM. * P < 0.05, *** P < 0.001 by 2-tailed t test. (D and F) Images of representative MCF7P or T47D xenograft tumors collected at the end points of the experiments in panel c or e, showing the manipulation of both GATA3 and JUN promotes stronger tumor *in vivo* growth than single gene manipulation. (G) A proposed model of coordinate role of GATA3 and AP1 in regulating enhancer reprogramming. Enhancer reprogramming mediated by the altered interactions between ERα and TFs (GATA3 and AP1) promote phenotypic plasticity during the acquisition of therapy resistance and invasive progression. In endocrine sensitive cells, ERα is recruited to sites with ERE motif and sites with GATA3 motif through *cis* and *trans* binding respectively to orchestrate an epithelial-like gene expression program. During the transition from endocrine sensitive to resistant phenotypes, loss of GATA3 expression leads to the loss of enhancers co-occupied by ERα and GATA3. Meanwhile, increased AP1 expression and activity turns on AP1 and ERα co-bound *GAIN* enhancers, resulting in an invasive cancer gene expression program.

To investigate the combined effect of AP1 and GATA3 on tamoxifen response *in vivo*, we generated orthotopic xenograft tumors using MCF7P cells with JUN OE or GATA3 KD. Compared to control, JUN OE resulted in a more rapid mammary tumor growth in the presence of tamoxifen. However, we did not detect any tumor growth from grafted MCF7P cells expressing GATA3 shRNA. This could be due to a broad regulatory role of GATA3 in controlling cell growth. Interestingly, GATA3 silencing in the background of JUN OE did not inhibit grafted tumor growth, moreover, it enhanced the effects of JUN OE in promoting tumor growth (Figures 7C and 7D), suggesting that JUN OE could somehow compensate for the loss of GATA3. We further confirmed the combined effect of AP1 and GATA3 on the growth of *in vivo* xenografts derived from T47D cells with stably integrated Dox-inducible JUN OE construct and/or GATA3 shRNA. While GATA3 KD alone inhibited tumor growth in the presence of tamoxifen, it also significantly enhanced the growth-promoting effects of JUN OE (Figures 7E and 7F). Using RNA-seq data of 34 different cancer types from TCGA database, we performed GSEA analyses on cancer hallmark gene sets. We found that high expression level of JUN was positively associated with the enrichment of EMT pathway in breast cancer, however high expression level of GATA3 was negatively correlated with EMT pathway in breast cancer (Figures S7C and S7D), indicating their opposite roles in EMT-related cancer progression *in vivo* and supporting that the combined effect of JUN OE and GATA3 KD surpass the effect of individual manipulation. Taken together, our *in vitro* and *in vivo* cancer growth data support that during the transition from tamoxifen-sensitive (MCF7P) to tamoxifen-resistant (TamR) phenotypes, loss of GATA3 and elevation of AP1 level and activity combine in their effect on promoting phenotypic plasticity, resulting in endocrine resistance and a more aggressive cancer phenotype.

## DISCUSSION

Therapy resistance is a life-threatening problem in cancer treatment and it often associates with cancer invasive progression. Phenotypic plasticity, which can be enhanced by epigenetic reprogramming, is recognized as a general cause of therapy resistance in many cancers including lung cancer, prostate cancer, and others ^3–6^. In ER+ luminal breast cancer, cell phenotypic transition during cancer progression is associated with the unsuccessful applications of tamoxifen and other endocrine agents that target ERα signaling ^48^. To understand the underlying mechanism of endocrine resistance, we take advantage of a cellular model of endocrine resistance (TamR) that derive from MCF7P breast cancer cell line. Transcriptional profiling revealed an upregulation of basal/mesenchymal and EMT signature genes, as well as a downregulation of luminal/epithelial markers in TamR cells, linking the development of tamoxifen resistance with the regulation of phenotypic plasticity. Further studies showed that the differential gene expression was driven by a wide spread reorganization of ERα enhancer landscape, which was mediated predominantly by GATA3 and AP1. In addition, manipulating GATA3 or AP1 expression was sufficient to alter cancer cell properties including morphology and sensitivity to endocrine therapy agents and these findings were extended to additional breast cancer cell lines as well as *in vivo* xenograft models. Therefore, we propose that a global enhancer gain/loss reprogramming driven by changes in TF interactions, particularly between ERα and GATA3 or AP1, profoundly alter breast cancer transcriptional programs to promote cellular plasticity and therapy resistance (Figure 7G). This is also supported by clinical data, in which we found strong correlation between cancer invasiveness and the expression level or activity of these two TFs and their targets.

Phenotypic and cellular plasticity are related to cancer progression and therapy response ^3^. The EMT process including a hybrid epithelial/mesenchymal cell state, in which cells acquire plasticity and gain stem cell-like properties, is the most widely studied example of plasticity in tumor progression ^3, 27^. During EMT, cells of a differentiated epithelial phenotype, which are polarized and form extensive cell-cell adhesions, lose their apicobasal polarity and become motile, associated with the expression of mesenchymal marker ^49^. During the transition from MCF7P to TamR, we observed morphological changes similar to those in EMT. We identified upregulated genes that are highly enriched for EMT associated signature genes and mesenchymal marker genes, as well as downregulated genes enriched for epithelial marker genes. We also found TamR cells were at a hybrid epithelial/mesenchymal cell state (Figure 1G), which is often found to associate with invasiveness and therapy resistance ^27^. Furthermore, this transition of gene expression positively correlated with enhancer landscape reorganization mediated by the differential interactions between ERα and other TFs, especially GATA3/AP1. Such transitions in cellular properties and in gene expression programs during the development of tamoxifen resistance bear an obvious commonality with those in EMT, suggesting that enhancer reprogramming promotes endocrine resistance via inducing plasticity.

Enhancers are bound and regulated by a mixture of common and lineage-specific TFs and cofactors to achieve cell-specific or context-specific transcription regulation. Similar to other enhancers, ERα enhancers are associated with various co-regulators. Using an *in vivo* biotin tagging system coupled with omics analysis, we have previously identified multiple oncogenic TFs (MegaTrans TFs) as ERα cofactors to determine enhancer activity ^19^. These MegaTrans TFs, besides binding to their conventional DNA binding motifs (in *cis* direct DNA binding), are also recruited by ERα through protein-protein interactions (in *trans* tethering-binding) to active ERE enhancers as ‘co-activators’. In this study, taking advantage of a powerful live-cell proximity biotin labeling BioID technology and other omics assays, we observed locus-specific binding pattern for ERα and many of its TF cofactors including FOXA1, GATA3 and AP1. On the *COMMON* enhancers between MCF7P and TamR cells, ERα binding was in *cis*, while GATA3 and AP1 appeared to bind to ERα enhancers in *trans* through tethering to ERα. In contrast, on the *LOSS* enhancers, occupancy of ERα was in *trans* and mediated by interacting with GATA3 that directly binds to GATA3 DNA motif on chromatin. Similarly, ERα was recruited to *GAIN* enhancers by *cis*-bound AP1 through protein-protein interaction. As expected, FOXA1 binding was found in all three groups of enhancers (*COMMON, LOSS* and *GAIN*). In addition to GATA3, AP1, and FOXA1, we identified motifs for other TFs including AP2γ, RUNX2 (Figures 3A and 3B), indicating that endocrine resistance may involve other TFs than GATA3, AP1, and FOXA1. This notion has been supported by our recent published findings that RUNX2-ERα interaction regulates a group of invasive genes in TamR cells ^22^. Thus, more studies are needed in the near future to functional tests for the other TFs. Collectively, our findings suggest that enhancer-TF interaction could be very complex, resulting in tightly controlled gene expression program during cancer progression.

Among the ERα-interacting TF cofactors, FOXA1 is a pioneer factor that functions in chromatin remodeling and accessibility. FOXA1 binds to its own DNA motifs adjacent to ERE to open condensed chromatin to facilitate ERα binding on enhancers, followed by the recruitment of epigenetic cofactors including histone acetyltransferase P300 and mediator MED1. Previous work has shown cooperativity between FOXA1 and ERα in controlling estrogen-induced gene expression ^50^. High levels of expression of FOXA1 and ERα have been found in breast cancer metastases that are resistant to endocrine therapy ^14^. Moreover, FOXA1 motif was found enriched in ERα binding sites that are specifically associated with patient’s relapse ^14^, connecting FOXA1 function with cancer invasiveness. More recently, our effort to characterize breast cancer endocrine resistant cell models discovered gene amplification of FOXA1 in two independent cell lines ^21^. FOXA1 overexpression at both mRNA and protein levels was also found in other tamoxifen resistant cell lines without FOXA1 amplification. Elevation of FOXA1 level in ER+ breast cancer cells activated multiple oncogenic pathways including IL8 signaling, resulting in endocrine resistance and increased cell invasion ^21, 51, 52^. In the current study, we found that FOXA1 binding was maintained in all three groups of enhancers, consistent with its role as a pioneer factor. Given the critical function of FOXA1 in regulating ERα enhancer activity during the development of endocrine resistance, increased FOXA1 levels through amplification might play an essential role in regulating both enhancer loss and gain reprogramming (together with GATA3 for *LOSS* enhancers and together with AP1 for *GAIN* enhancers). Consistent with this notion, FOXA1-dependent reorganization of enhancer landscape can promote pancreatic cancer metastasis ^53^. Therefore, FOXA1 might function with other TFs to direct enhancer reprogramming to promote cancer progression in different tumors.

This study for the first time identified a coordinate role of GATA3 and AP1 in mediating enhancer reprogramming. Our data showed that GATA3 and AP1 each regulate a different gene program: GATA3 controls the luminal lineage-specific gene program and AP1 regulates cancer invasion-related gene program including basal/mesenchymal marker genes (Figure 7G). Interestingly, although GATA3 did not bind to *GAIN* enhancers, GATA3 KD appeared to synergize the effect of JUN OE on *GAIN* enhancers activation, suggesting that loss of GATA3 might indirectly influence JUN-mediated enhancer regulation through an unknown mechanism. One potential explanation is that GATA3 may compete with other TFs like AP1 to recruit FOXA1 or chromatin remodeling complex on chromatin. When GATA3 is decreased or lost, FOXA1 or chromatin remodeling complex can be released to cooperate with AP1 to bind at *GAIN* enhancers, which is reminiscent of two previous reports that FOXA1 downregulation can relinquish AR to permissively bind AREs across the genome ^54, 55^. Given the coordinate role of GATA3 and AP1 in regulating enhancers, we expected that simultaneously knocking down GATA3 and overexpressing JUN would have synergistic effect on promoting cancer growth in the presence of tamoxifen. However, we only detected additive effect in the *in vitro* cell growth assay. It was also not expected that knocking down GATA3 completely inhibited xenograft tumor growth but enhanced tumor growth in the background of JUN overexpression. These results suggest a broad gene regulation role for each TF and a complex functional interaction between GATA3 and AP1 in ER+ tumor progression.

Collectively, the central conclusion of this study is that altered interactions between ERα and TF cofactors GATA3 and AP1 lead to enhancer reprogramming that promotes plasticity and endocrine resistance. However future studies are required to address many unanswered questions including: 1) How does GATA3 maintain *LOSS* enhancers? 2) How does AP1 activate *de novo GAIN* enhancers? 3) How do different TFs (e.g. ERα, FOXA1, GATA3, AP1 and even more others) coordinate to control enhancer activity? 4) What are the microenvironmental signals that trigger changes in expression and/or activity of GATA3 and AP1 during the acquisition of hormone resistance?

## ACKNOWLEDGMENTS

This work was supported by grants from CPRIT award (RR160017 to Z.L.), V Foundation’s V Scholar Award (V2016-017 to Z.L.) and V Scholar Plus Award (DVP2019-018 to Z.L.), Max and Minnie Tomerlin Voelcker Fund Young Investigator Award (to Z.L.), Susan G. Komen CCR award (CCR17483391 to Z.L.), and National Cancer Institute (U54 CA217297/PRJ001 to Z.L.), and UT Rising STARs Award to Z.L.. All deep sequencing data was generated in the Genome Sequencing Facility at Greehey Children’s Cancer Research Institute in the UT Health San Antonio, which is supported by NIH-NCI P30 CA054174 (NIH Cancer Center at UT Health San Antonio), NIH Shared Instrument grant 1S10OD021805-01 (S10 grant), and CPRIT Core Facility Award (RP160732). BASiC, where pyrosequencing assay experiments were carried out, was supported by the CPRIT award (RP150600) and funding from the Office of Vice President of Research, UTHSCSA. R.S. and X.F. were supported by the NIH SPORE Grants (P50 CA058183 and CA186784 to R.S.), Cancer Center Grants (P30 CA125123 and P30 CA008748 to R.S. and X.F.), Breast Cancer Research Foundation (BCRF-17-143 and 18-145 to R.S.), CPRIT grant RP190398 (R.S. and X.F.) and DOD grant (W81XWH-14-1-0326 to X.F.).

## AUTHOR CONTRIBUTIONS

M.B. performed most of the experiments with assistance from H.W., P.X. and Z.L.. Z.Z. performed the computational analyses for all next-generation sequencing assays. Z.L., M.B. and Z.Z. conceived the work and designed the study. Z.L. supervised the research and oversaw the project. Z.L., L.C., M.B., and Z.Z. wrote the manuscript with input from all authors. E.M. provided paired parental vs tamoxifen-resistant human clinical PDX samples and microarray data. X.F., C.A. and R.S. provided paired tamoxifen-sensitive and -resistant MCF7 cells for this study, provided comments, and reviewed the manuscript.

## ADDITIONAL INFORMATION

Supplementary information is available for this paper (including 3 Excel tables, 7 supplemental figures and their figure legends).

## METHODS

### Cell culture studies

T47D cell line was obtained from ATCC. T47D cells were cultured in Roswell Park Memorial Institute (RPMI) 1640 Medium supplemented with 10% FBS and penicillin/streptomycin. Paired tamoxifen-sensitive and -resistant MCF7 cells were provided by Dr. Rachel Schiff at Baylor College of Medicine, Houston, TX. Authenticity of each cell line was confirmed once the resistance to tamoxifen was established. MCF7 parental cells were maintained in RPMI 1640 supplemented with 10% heat-inactivated FBS (Sigma) and 1% penicillin-streptomycin (P/S). The endocrine-resistant cells were kept in phenol-red free medium supplemented with 10% heat-inactivated charcoal-stripped-FBS and 1% P/S with the addition of 100 nM 4-hydroxytamoxifen (4-OHT, H7904, Sigma). All cells were kept at 37°C in a humified incubator with 5% CO_2_.

### Generation of doxycycline-inducible MCF7P and TamR stable cell lines

To set up the doxycycline-inducible GATA3 and JUN-overexpressing cell lines, GATA3 and JUN cDNAs (Open Biosystems) were individually cloned into pCR8/GW/TOPO and then transferred to the pInducer20 destination vector (pInducer20 was a gift from Stephen Elledge; Addgene #44012) using the Gateway system (Invitrogen), followed by co-transfection with packaging plasmids (psPAX2 and pMD2.G from Addgene) into 293T cells. Culture medium containing lentivirus particles were harvested, filtered, and used to infect cells in media containing polybrene (8 µg/ml). Cells were selected by 500 µg/ml G418 (Invitrogen) after infection to set up doxycycline-inducible stable cell lines.

### shRNA lentivirus package and infection

Mission shRNA lentiviral plasmids targeting GATA3, JUN and control shRNA were purchased from Sigma (their catalog numbers are listed in Supplementary Table S1). Knockdown experiments with shRNA lentiviruses were conducted according to the standard lentivirus package and transduction protocols from Addgene. These pLKO-based lentiviral shRNA plasmids were co-transfected with packaging plasmids (psPAX2 and pMD2.G) into 293T cells. Culture medium containing lentivirus particles were harvested, filtered, and used for cell infection. Stable knockdown cells were selected with 1 µg/ml puromycin and collected for experiments within 3-5 days.

To set up doxycycline inducible GATA3 knockdown cell line, GATA3 shRNA was sub-cloned into Tet-pLKO-puro, a gift from Dmitri Wiederschain (Addgene #21915). Stable cell lines were generated after puromycin selection. Doxycycline at a concentration of 100 ng/ml was used to achieve GATA3 knockdown.

### Generation of BLRP-tagged inducible MCF7P and TamR stable cell lines and biotin ChIP-qPCRs

To study binding patterns of wildtype and DNA-binding domain mutant (pBox mutant) of ERα in MCF7P or TamR cell line, we used an *in vivo* biotinylation tagging BirA-BLRP system as previously described ^19^. The BLRP-tagged wild-type and pBox mutant ERα cDNAs were cloned into a retrovirus-based Tet-On expression vector pRetroX-Tight-Pur (Clontech #632104) ^19^. These constructs were co-transfected with pCL-Ampho packaging plasmid into 293T cells for retrovirus production. Then, the retroviruses were transduced into parental MCF7P and TamR stable cell lines that have been engineered to express BirA enzyme and Tet Repressor. G418 (500 µg/ml), hygromycin (200 µg/ml), and puromycin (0.5 µg/ml) were used for selection and stable cell line maintenance. To induce BLRP-tagged protein expression, stable cell lines were treated with 2 µg/ml doxycycline for approximately 24 hours and were collected to check for BLRP-tagged protein expression levels by immunoblotting with specific antibodies before biotin ChIP experiments.

Biotin ChIP-seq experiments for BLRP-tagged wildtype or mutant ERα were performed with our previous protocol^19^. Briefly, cross-linked protein-DNA complexes were pulled down by MyOne Streptavidin T1 Dynabeads (Thermo-Fisher Scientific) and the washing was performed under much more stringent conditions that included 4 washes with 1% SDS in TE (20 min each) and two washes with 1% Triton X-100 in TE. The washed streptavidin beads were then subjected to TEV protease (Life Technologies) digestion twice for tagged protein-DNA complex elution before de-crosslinking at 65°C overnight. The following day, the final ChIP DNA was purified and resuspended in 10 mM Tris-HCl pH 8.5. The purified DNA was subjected to qPCR directly to confirm target region enrichment. BLRP-tagged ERα stable cell lines without doxycycline induction were used as controls for background detection. All primers used for qPCR are listed in Table S1.

### RNA isolation and quantitative RT-PCR

Total RNA was isolated with RNeasy Mini Kit (Qiagen) according to the manufacture’s protocol and 1 µg RNA was used to convert to cDNA using iScript Select cDNA Synthesis Kit (Bio-Rad) in the presence of both oligo (dT) and random primers. qPCR was conducted with SsoAdvanced Universal SYBR Green Supermix (Bio-Rad) using CFX384 Real-Time PCR Detection System (Bio-Rad) according to the manufacturer’s instructions. Relative expression of RNAs was determined by the ΔΔCT method using GAPDH or ACTB as an internal control for quantification analyses of gene targets. The primers used for qPCR are listed in Table S1.

### RNA-seq

Total RNA was extracted using RNeasy Mini Kit (Qiagen). For each sample, 1 µg RNA was used for library construction using KAPA RNA HyperPrep Kit with RiboErase (KK8560) or KAPA mRNA HyperPrep Kit (KK8580). For sequencing, samples with specific adaptors were sequenced with the Illumina’s HiSeq 3000 system according to the manufacturer’s instructions.

### Western blot

Cells were lysed in RIPA lysis buffer (50 mM Tris-Cl pH 8.0, 150 mM NaCl, NP-40, 0.5% sodium deoxycholate, 0.1% SDS) supplemented with 1 mM DTT, 1 mM PMSF, and 1x protease inhibitor cocktail (Roche). Protein concentrations were quantified with the Bio-Rad protein assay kit. Western blotting was performed as previously described^19^. Briefly, 30 µg of protein extracts were loaded and separated by SDS–PAGE gels. Blotting was performed with standard protocols using PVDF membrane (Bio-Rad). Membranes were blocked for 1 hour in blocking buffer (5% Non-fat milk in PBST) and probed with primary antibodies at 4°C overnight. After three washes with PBST, the membranes were incubated with HRP-conjugated secondary antibody. Signals were visualized with Clarity Western ECL Substrate (Bio-Rad) as described by the manufacturer. All antibodies used for immunoblotting are listed in Table S2.

### Co-immunoprecipitation

MCF7P or TamR cellular extracts were prepared by incubating cells with NP-40 lysis buffer (50 mM Tris-HCl pH 8.0, 150 mM NaCl, 0.5% NP-40, 1 mM PMSF and 1x protease inhibitor) for 30 min on ice. Supernatants were collected by centrifugation at 13,000rpm for 15 min at 4°C. For immunoprecipitation, 500 µg of protein was incubated with 3 µg of ERα antibody or 3 µg of mouse IgG overnight at 4°C with rotation. 20 µl of Dynabeads Protein G (Invitrogen) was then added and incubated for an additional 2 hours. Then, the bead-protein complexes were washed five times with NP-40 lysis buffer. The precipitated proteins were eluted from the beads with 2x SDS loading buffer and boiling for 10 min followed by western blot analyses. The antibodies used for co-IP are listed in Table S2.

### Immunofluorescence staining

Cells were grown on poly-D-lysine–coated coverslips for 24 hours. After washing twice with PBS, cells were fixed in 4% paraformaldehyde for 10 min and permeabilized by 0.1% Triton X-100 in PBS for 10 min at room temperature. Then, the cells were blocked using 1% BSA/PBS for 1 hour and sequentially incubated with the primary antibody in 0.1% BSA/PBS at 4°C overnight. After washing, immunoreactivity was developed using anti-mouse or rabbit IgG conjugated with Alexa-Fluor-488 (Jackson ImmunoResearch). Coverslips were mounted by mounting medium with DAPI (Life Technologies) and sealed with nail polish before imaging with Cytation 5 (BioTek). The antibodies used for immunofluorescence staining are listed in Table S2.

### BioID system setup and pulldown experiment

The pcDNA3.1 MCS-BirA(R118G)-HA (Addgene plasmid #36047) were kindly provided by Kyle Roux ^33^. To make the tet-on inducible BioID constructs, BirA(R118G)-HA from pcDNA3.1 MCS-BirA(R118G)-HA plasmid was cloned into pRetroX-Tight-Pur (Clontech) at EcoRI site by PCR to get pRetroX-MCS-BioID-HA vector. The full-length human ERα cDNA was then cloned into pRetroX-MCS-BioID-HA at the BamHI and MluI sites to get pRetroX-ERα-BioID-HA. The pRetroX-ERα-BioID-HA was co-transfected with pCL-Ampho packaging plasmid into 293T cell line to produce retrovirus. The retrovirus was transduced into parental MCF7P and TamR cell lines that were engineered to stably express a Tet Repressor using a retroviral vector. Stable cell lines were selected using 200 µg/ml hygromycin and 0.5 µg/ml puromycin. To induce ERα-BioIDHA protein expression, 2 µg/ml doxycycline was added into culture media approximately 24 hours before adding 50 mM Biotin for another 24 hours to label all proteins in the proximity of ERα before collection.

To detect ERα-associated cofactors on chromatin, we used nuclear fraction for biotin-labeled proteins. Cytoplasmic and nuclear lysates were isolated as previously described with slight modification ^56^. Briefly, cells were scraped from 15cm dish plates and washed with cold PBS. For every 70 mg of cells, 1 ml of cold Hypotonic Lysis Buffer (HLB) (10 mM Tris-HCl pH 7.5, 3 mM MgCl_2_, 10 mM NaCl, 0.3% NP-40, 10% glycerol and 1x protease inhibitor) was used to resuspend the cells and the cell suspension was then incubated on ice for 10 min followed by centrifugation at 800g for 8 min at 4°C. The supernatant was collected as cytoplasmic lysate fraction. The precipitated nuclei were washed 4 times in HLB with pipetting and centrifuging at 200g for 2 min at 4°C. After HLB washes, the nuclei were resuspended in 0.5 ml of ice-cold Modified RIPA buffer (MRB) (50 mM Tris-HCl pH 7.4, 0.1% SDS, 0.5% Sodium Deoxycholate, 1% NP-40, 6 mM MgCl_2_, 150 mM NaCl, 1 mM EDTA, 1 mM DTT, and 1x protease inhibitor). The nuclei pellet in MRB was subjected to sonication till the solution turned clear. The solution was then treated with 25 U/ml Benzonase and 4 U/ml DNaseI for 30 min at RT to digest the chromatin, followed by centrifugation at 18,000g for 15 min at 4°C to pellet insoluble protein and other components. The supernatant was collected as nuclear lysate fraction for the pulldown experiments. For each sample, a small aliquot for each fraction was used to test the cellular fractionation efficiency with GAPDH and Histone H3 western blots before proceeding with BioID pulldown experiments below.

For BioID pulldown, each nuclear lysate fraction was incubated with 200 µl PBS-washed MyOne Streptavidin T1 Dynabeads (Life Technologies) with overnight rotation at 4⁰C to pull down biotinylated proteins in nuclear lysate. The beads were then washed for five times with RIPA buffer (50 mM Tris PH 7.4, 0.4% SDS, 0.1% Triton X-100, 1% NP-40, 300 mM NaCl, 1 mM EDTA, 0.5% sodium deoxycholate, 1 mM DTT) with 10 min rotation for each wash (Note: only the first two washes need to have protease inhibitor and incubated at 4⁰C. The last three washes can be done at RT without protease inhibitor). After 5X washes with RIPA, the beads were washed 2X with stringent RIPA buffer containing 5% SDS, followed by 4X washes with PBS to get rid of SDS residue that might affect trypsin digestion during mass spectrometry, with only 200 µl PBS used for the 4^th^ wash. 10% (20 µl) of the beads suspension was transferred to another tube for western blot and 90% (180 µl) was kept in the original tube for mass spectrometry. Both tubes were placed on a magnetic stand for 3 min to remove PBS completely. For western blot, the beads were boiled for 10 min in 2X protein loading buffer with 5% β-Me to release proteins. For mass spectrometry, beads were stored at -80⁰C before shipping to the Proteomics Core at Sanford-Burnham-Prebys Medical Discovery Institute.

### Mass spectrometry for BioID pulldown beads

Following immunoprecipitation and washes, proteins were digested directly on-beads. Proteins bound to the beads were resuspended with 8 M urea, 50 mM ammonium bicarbonate, and cysteine disulfide bonds were reduced with 5 mM tris(2-carboxyethyl)phosphine (TCEP) at 30°C for 60 min followed by cysteine alkylation with 15 mM iodoacetamide (IAA) in the dark at room temperature for 30 min. Following alkylation, urea was diluted to 1 M using 50 mM ammonium bicarbonate, and proteins were finally subjected to overnight digestion with mass spec grade Trypsin/Lys-C mix (Promega, Madison, WI). Finally, beads were pulled down and the solution with peptides collected into a new tube. The beads were then washed once with 50 mM ammonium bicarbonate to increase peptide recovery. Following overnight digestion, samples were acidified with formic acid (FA) and subsequently desalted using AssayMap C18 cartridges mounted on an Agilent AssayMap BRAVO liquid handling system. C18 cartridges were first conditioned with 100% acetonitrile (ACN), followed by 0.1% FA. Samples were then loaded onto the conditioned C18 cartridge, washed with 0.1% FA, and eluted with 60% ACN, 0.1% FA. Finally, the organic solvent was removed in a SpeedVac concentrator prior to LC-MS/MS analysis. Before injecting in the LC-MS, total sample peptide amount was determined by Pierce Quantitative Colorimetric Peptide Assay, 500 Assays (ThermoFisher).

Dried samples were reconstituted with 2% acetonitrile, 0.1% formic acid and analyzed by LC-MS/MS using a Proxeon EASY nanoLC system (Thermo Fisher Scientific) coupled to Elite mass spectrometer (Thermo Fisher Scientific). Peptides were separated using an analytical C18 Acclaim PepMap column 75µm x 500mm, 2µm particles (Thermo Scientific) in 121 min at a flow rate of 300 nl/min: 1% to 6% B in 1 min, 6% to 23% B in 56 min, 23% to 34% B in 37 min, 34% to 48% B in 26 min, and 48% to 98% B in 1 min (A = Formic acid 0.1%; B = 80% ACN: 0.1% Formic acid).

The mass spectrometer was operated in positive data-dependent acquisition mode. MS1 spectra were measured in the Orbitrap with a resolution of 60,000 (AGC target of 1e6, and a mass range from 350 to 1450 m/z). Up to 10 MS2 spectra per duty cycle were triggered, CID-fragmented and acquired in the linear ion trap (AGC target of 1e4, isolation window of 2m/z, and a normalized collision energy of 35). Dynamic exclusion was enabled with a duration of 30 seconds.

### Proteomics data analysis

MS raw files were analyzed by MaxQuant software with most default settings, and Andromeda search engine was used to search against the human Uniprot database. False discovery rate was set to 0.01 for proteins and peptides with a minimum length of seven amino acids. Variable modifications were set to methionine oxidation and N-terminal acetylation; while fixed modification was set to carbamidomethylation. For label-free protein quantification ^57^, the minimum ratio count was set to two, and peptides for quantification were set to unique and razor. Statistical analysis was performed by Perseus ^58^. Proteins identified only by site, reverse hits or potential contaminants were removed before downstream analysis. LFQ intensities were log2 transformed, and the matrix was then grouped with regard to cell lines. The proteins were then filtered by requiring at least two valid values in at least one cell line. Missing values were imputed based on normal distribution. Paired t-test was then used to determine the significant difference of protein level between the two cell lines.

### ChIP-seq

ChIP assays were performed as previously described ^19^ with slight modification. Briefly, cells were cross-linked with 1% formaldehyde for 10 min at room temperature. For selected experiments (e.g., ChIP for P300, JUN, BRG1 and ARID1B), cells were double cross-linked with 2 mM DSG (CovaChem) for 45 min followed by secondary fixation with 1% formaldehyde for 10 min. Cross-linking was quenched with 0.125 M glycine for 5 min. Cells were successively lysed in lysis buffer LB1 (50 mM HEPES-KOH, pH 7.5, 140 mM NaCl, 1 mM EDTA, 10% glycerol, 0.5% NP-40, 0.25% Triton X-100, 1x PI), LB2 (10 mM Tris-HCl, pH 8.0, 200 mM NaCl, 1 mM EDTA, 0.5 mM EGTA, 1x PI), LB3 (10 mM Tris-HCl, pH 8.0, 100 mM NaCl, 1 mM EDTA, 0.5 mM EGTA, 0.1% Na-Deoxycholate, 0.5% N-lauroylsarcosine, 1x PI) (Note: after passing through buffers LB1 and LB2, the pellet becomes nuclear fraction and will be lysed in LB3 lysis buffer for sonication). Chromatin was sonicated to an average size of ∼200-500 bp using QSonica’s Q800R sonicator system (20% amplitude, 10s on and 20s off for 10 min). A total of 3-6 µg of antibody was added to the sonicated chromatin and incubated overnight at 4℃. Subsequently, 50 µl of Dynabeads Protein G (Invitrogen) were added to each ChIP reaction and incubated for 4 hours at 4℃. Dynabeads were washed with RIPA buffer (50 mM HEPES pH 7.6, 1 mM EDTA, 0.7% Na-Deoxycholate, 1% NP-40, 0.5 M LiCl) 6 times, and once with TE. The chromatin was eluted, followed by reverse cross-linking and DNA purification. ChIP DNA was resuspended in 10 mM Tris-HCl pH 8.5. All antibodies used in this study are summarized in the Table S2. The purified DNA was subjected to qPCR to confirm target region enrichment before moving on to deep sequencing library preparation. For sequencing, the extracted DNA was used to construct the ChIP-seq library using KAPA Hyper Prep kit (KK8504), followed by deep sequencing with the Illumina’s HiSeq 3000 system according to the manufacturer’s instructions.

### ATAC-seq

ATAC-seq library prep was performed as previously described^59^. Briefly, 50,000 cells were washed three times with cold PBS, collected by centrifugation then lysed in lysis buffer (10 mM Tris-HCl, pH 7.4, 10 mM NaCl, 3 mM MgCl2, 0.1% NP-40). After purification of nuclei, transposition was performed with Tn5 transposase from Nextera DNA Library Prep Kit (Illumina, catalog # FC-121-1030). Purified DNA was then ligated with adapters, amplified and size selected for sequencing. Libraries were sequenced with Mid 75 bp PE on Illumina NextSeq 500 (4 samples in one lane).

### Global run-on sequencing (GRO-seq)

GRO-seq experiments were performed as previously described ^18, 19^. MCF7P or TamR cells growing in 15cm plate with full media were washed three times with cold PBS and then sequentially swelled in swelling buffer (10 mM Tris-HCl pH7.5, 2 mM MgCl_2_, 3 mM CaCl_2_) for 10 min on ice, harvested, and lysed in lysis buffer (swelling buffer plus 0.5% NP-40, 20 units of SUPERase-In, and 10% glycerol). The resultant nuclei were washed two more times with 10ml lysis buffer and finally re-suspended in 100 µl of freezing buffer (50 mM Tris-HCl pH 8.3, 40% glycerol, 5 mM MgCl_2_, 0.1 mM EDTA). For the run-on assay, re-suspended nuclei were mixed with an equal volume of reaction buffer (10 mM Tris-HCl pH 8.0, 5 mM MgCl_2_, 1 mM DTT, 300 mM KCl, 20 units of SUPERase-In, 1% sarkosyl, 500 µM ATP, GTP, and Br-UTP, 2 µM CTP) and incubated for 5 min at 30°C. The resultant nuclear-run-on RNA (NRO-RNA) was then extracted with TRIzol LS reagent (Life Technologies) following manufacturer’s instructions. NRO-RNA was fragmented to ∼300-500 nt by alkaline base hydrolysis on ice for 30 min followed by treatment with DNase I and antarctic phosphatase. At this step, only a small portion of all the RNA species were BrdU-labeled. To purify the Br-UTP labeled nascent RNA, the fragmented NRO-RNA was immunoprecipitated with anti-BrdU argarose beads (Santa Cruz Biotechnology) in binding buffer (0.5XSSPE, 1 mM EDTA, 0.05% tween) for 1-3 hours at 4°C with rotation. Subsequently, T4 PNK was used to repair the ends of the immunoprecipitated Br-UTP labeled nascent RNA at 37°C for 1 hour. The RNA was extracted and precipitated using acidic phenol-chloroform.

The RNA fragments were subjected to poly-A tailing reaction by poly-A polymerase (NEB) for 30 min at 37°C. Subsequently, reverse transcription was performed using oNTI223 primer and superscript III RT kit (Life Technologies). The cDNA products were separated on a 10% polyacrylamide TBE-urea gel and only those fragments migrating between 100-500 bp were excised and recovered by gel extraction. Next, the first-strand cDNA was circularized by CircLigase (Epicentre) and re-linearized by APE1 (NEB). Re-linearized single strand cDNA (sscDNA) was separated on a 10% polyacrylamide TBE gel and the appropriately sized product (∼120-320 bp) was excised and gel-extracted. Finally, sscDNA template was amplified by PCR using the Phusion High-Fidelity enzyme (NEB) according to the manufacturer’s instructions. The oligonucleotide primers oNTI200 and oNTI201 were used to generate DNA libraries for deep sequencing. The sequences for primers oNTI223, oNTI200 and oNTI201 are listed in Table S1.

### Pyrosequencing assay

500 ng of genomic DNA per sample was used for bisulfite conversion with the EZ DNA Methylation™ Kit (Zymo Research) according to the manufacturer’s instruction. PCR was performed with primers that flank CpG sites with differential methylation levels between MCF7P and TamR cells at the GATA3 promoter region. The biotinylated PCR product was then sequenced with a sequencing primer. The methylation levels were quantified by the PyroMark Q96 MD System at the Bioanalytics and Single-Cell Core (BASiC) of UTHSCSA. The methylation percentage of each interrogated CpG site was calculated by PyroMark CpG software (Qiagen, Carlsbad, CA). Details of primer sequence for pyrosequencing assay are listed in Table S1.

### Cell proliferation assays

Cell proliferation was assessed using Cell Counting Kit-8 (CCK-8, Sigma) following manufacturer’s instructions. 2,000 cells/well were seeded in a 96-well plate. After 24 hours, cells were treated with different concentration of 4-OHT. The plates were pre-incubated in a humidified incubator at 37°C for 7 days, and 10 µl of the CCK-8 solution was then added to each well. The plate was incubated for 1-4 hours at 37°C. The absorbance was measured at 450 nm using a Cytation™ 5 imaging reader (Biotek).

### *In vivo* xenograft experiments

5 to 6-week-old athymic nude female mice (The Jackson Laboratory #002019) were housed in a 12 hours light/dark cycle in the animal facility at the UTHSCSA. All xenograft experiments were performed in accordance with a protocol approved by the Institutional Animal Care and Use Committee (IACUC) at the UTHSCSA. The study is compliant with all relevant ethical regulations regarding animal research. Estrogen pellets (Innovative Research, 0.72 mg 60-day release) were implanted underneath the skin on back three days prior to cell implantation. For orthotopic xenograft studies, 5×10^6^ MCF7P or T47D cells resuspended in 100 µl of 1:1 mix of growth media and Matrigel were injected into the fourth mammary fat pads of the mice using 27G 1/2 inch 1-ml syringe. For doxycycline induction experiments, mice were randomly placed in the treatment group or control group. 200 mg/L doxycycline was supplemented in their daily drinking water containing 0.5% sucrose. Tamoxifen was injected subcutaneously at a dose of 1 mg per mouse three times per week. Tumors were measured with Vernier calipers once a week and the tumor volumes were calculated as Volume (mm^3^) = π×length×width×height/6.

### DNA methylation data and TCGA data analysis

Illumina’s HumanMethylation450K BeadChip data for different cell lines including MCF7, TamR, MCF7X and FASR were retrieved from GEO website (GSE69118), with two replicates for each cell line ^39^. The raw data was preprocessed, and the background was normalized with the Bioconductor package minfi as described previously^60^. For visualization in IGV genome browser, a bedGraph file for each sample was generated using the genome position and beta value of each probe, and then transformed to bigwig file by bedGraphToBigWig command from UCSC^61^.

Genome-wide DNA methylome in TCGA was retrieved from Firehose ^62^. Only those samples that have both DNA methylation and mRNA-Seq data available were retained. Any methylation probes with missing data of more than 50 were excluded from analysis. All the patient samples were ordered based on the GATA3 expression level. The pearson correlation of GATA3 expression level with the beta value of each probe located 10kb upstream of the transcription start site (TSS) to the transcription end site (TES) was calculated. Breast cancer subtype information was also considered.

We retrieved the level 3 RNA-seq data for all 34 different cancer types in TCGA with data version 2016_01_28 from Firehose, and then calculated the correlation of GATA3 or JUN with all other genes across all tumor samples within each cancer type. After that, the genes were ranked from the highest positive correlation to lowest negative correlation, and GSEA analysis was performed for these ranked lists against cancer hallmark gene sets from MSigDb database. The gene sets were ranked based on the average normalized enrichment score (NES) across all cancer types.

### RNA-seq and microarray data analysis

The sequencing reads were aligned to human genome (hg19) with STAR 2.5.2b, a spliced-read aligner, and after removing reads mapped to rRNA sequences, read counts for each gene were conducted by featureCounts package with default parameter ^63, 64^. Genes with less than 1 read in at least 2 samples were discarded. DESeq2 1.14.1 was used to call differentially expressed genes with fold expression change >= 1.5 and FDR <= 0.05 as the cutoff ^65^. The log2 fold change from this analysis was then used to perform a pre-ranked GSEA analysis using JAVA GSEA 2.2.0 program by searching against the Broad Molecular Signatures Database (MsigDB) gene sets v5 ^24^. Moreover, we retrieved our published microarray dataset for the paired PDX models ^66^, and performed GSEA analyses on this dataset (Figure 1F).

### ChIP-seq data analysis

Reads were aligned to human genome (hg19) using bowtie with “--best --strata –m 1” parameters ^67^. Only uniquely mapped and non-duplicated reads were selected for subsequent analysis. MACS2 was employed to call peaks comparing immunoprecipitated chromatin with input chromatin using standard parameters and a q-value cutoff of 1e-5 ^68^. For H3K4me1 and H3K27ac, the broad mode of MACS2 was switched on. ROSE was used to identify super-enhancers based on the ranking of H3K27ac signal intensities, with stitching distance set to 15,000 bp and exclusion region distance to TSS set to 2,500 bp ^69^. The peaks overlapping with the blacklist regions from UCSC were removed. ChIPseeker was used to annotate the peaks ^70^. GREAT was used to annotate the potential functions of the peaks with default parameters ^28^.

For motif discovery, findMotifs.pl program in HOMER was used for the regions with 300 bp upstream and downstream of the peak summit ^71^. For motif distribution along the TamR-high to MCF7P-high enhancers that are defined by differential binding intensity of ERα in these two cell lines, FIMO was used to scan each enhancer from start to end ^72^. The region with the maximum motif score was assigned to that motif. If the p-value for this region was less than 1e-4, the enhancer will be considered containing the motif.

For integrative analysis of gene expression and ChIP-seq data, we first determined the closest gene for each enhancer, and then stratified the genes based on the enhancer number closest to the TSS site of each gene. For the net enhancer change in our analyses, total number of TamR-specific enhancers minus total number of MCF7P-specific enhancers was used.

### ATAC-seq data analysis

Cutadapt 1.11 was used to trim adapters in ATAC-seq reads, which were then aligned to human genome (hg19) using bowtie with parameters “--best --strata –m 1 –v 2”^73^. Aligned reads with the same genomic position and orientation were collapsed to a single read. The reads were extended to 200bp and normalized to a sequencing depth of ten million reads for each library.

### GRO-seq data analysis

GRO-seq reads were aligned to human genome (hg19) using bowtie with “--best --strata –m 1 –v 2” parameters. Duplicated reads were eliminated for subsequent analysis. To balance the clonal amplification bias and total useful reads, only three reads at most were allowed for each unique genomic position. When measuring the expression level of genes, mapped reads from the first 30 kb of gene body were counted, excluding promoter-proximal region (transcription start site (TSS) to 1000 bp downstream of TSS; if the length of a gene is shorter than 10 kb, then the reads mapped to the first 10%*length regions were excluded). If the length of a gene is shorter than 30 kb, then the mapped reads from the whole gene were counted, excluding promoter-proximal region and gene end (500 bp upstream of transcription termination site (TTS) to TTS). Differential expression analysis was then performed by DESeq2 with a fold change threshold of 2 and FDR<=0.01.

### Data Visualization

RNA-seq data was visualized in Integrative Genomics Viewer (IGV). Visualization of the data for ChIP-seq and GRO-seq was performed by organizing custom tracks onto the University of California, Santa Cruz, (UCSC) genome browser. ChIP-seq and GRO-seq samples were normalized to 10 million mapped reads per experiment, while RNA-seq samples were normalized to 1 million reads.

### Statistical Analysis

For all qPCRs and cell proliferation assays, statistical analyses were performed using two-tailed Student’s t test. The results were shown as mean ± SEM. Results are representative of at least three independent experiments. Significant differences between two groups were noted by asterisks (* P<0.05, **P<0.01, *** P<0.001).

### Data Availability

All deep sequencing raw data supporting the findings of this study have been deposited in the Gene Expression Omnibus (GEO) under accession code: GSE128460. All BioID proteomic data from MCF7P and TamR cell lines have been deposited in the Proteomexchange repository under the accession number PXD014015. Previous published DNA methylation data for different cell lines including MCF7, TamR, MCF7X and FASR were retrieved from GEO website (GSE69118). Previous published microarray dataset for paired PDX models including HBCx22 and HBCx22TamR were under GEO number GSE55561. All other data supporting the findings of this study are available from the corresponding author upon reasonable request.

## SUPPLEMENTARY FIGURE LEGENDS

**Figure S1.**
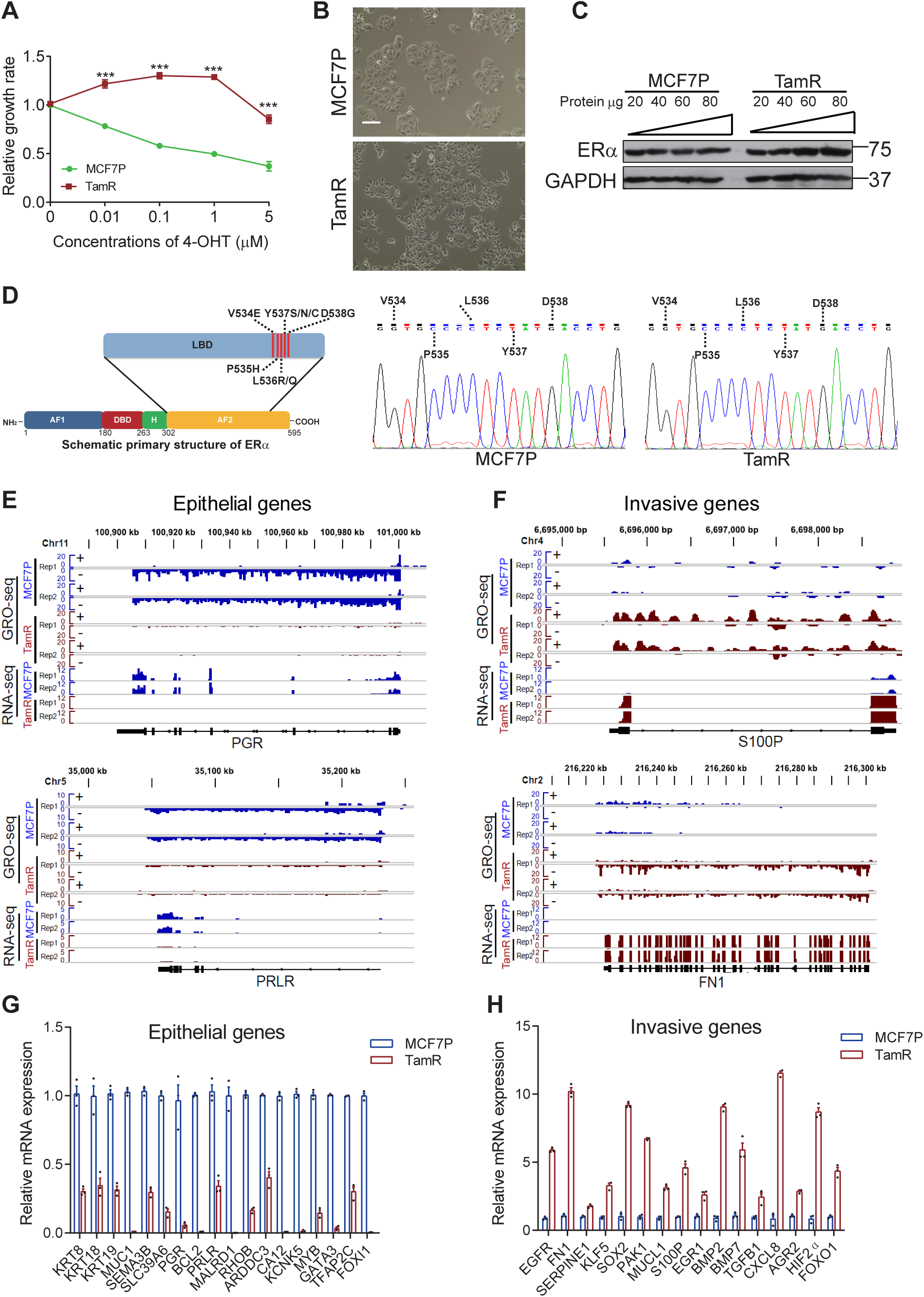
Endocrine resistance is associated with plasticity-enhancing transcriptome changes, Related to Figure 1. (A) The cell growth rate tests of MCF7P and TamR lines in the presence of 4-OHT showing the endocrine resistance of TamR line. (B) Brightfield images of MCF7P and TamR lines at ×100 magnification showing different morphology for these two lines. TamR cells morphologically mimic mesenchymal cells. Scale bar, 100 µm. (C) ERα protein levels in MCF7P and TamR cells detected by Western blots using a serial dilution of whole cell extract for semi quantitative purpose. GAPDH was used as a loading control. (D) Structural diagram of ERα protein showing the positions of point mutations in the ligand-binding domain (LBD) that were reported in endocrine-resistant or metastatic ERα+ breast cancers before (left). No LBD point mutation was detected in this TamR cell line with Sanger sequencing showing (right). (E) Genome browser snap images of the GRO-seq and RNA-seq signals at gene body regions for *PGR* and *PRLR* showing the significant downregulation of these two epithelial markers in TamR cells. (F) Genome browser snap images of the GRO-seq and RNA-seq signals at gene body regions for *S100P* and *FN1*, showing the significant upregulation of these two cancer invasiveness-associated genes in TamR cells. (G) RT-qPCR analysis of mRNA levels of selected epithelial markers in MCF7P and TamR cell lines. All of these epithelial markers are downregulated in TamR cells. (H) RT-qPCR analysis of mRNA levels of selected invasive genes in MCF7P and TamR cell lines. All of these invasive genes are upregulated in TamR cells.

**Figure S2.**
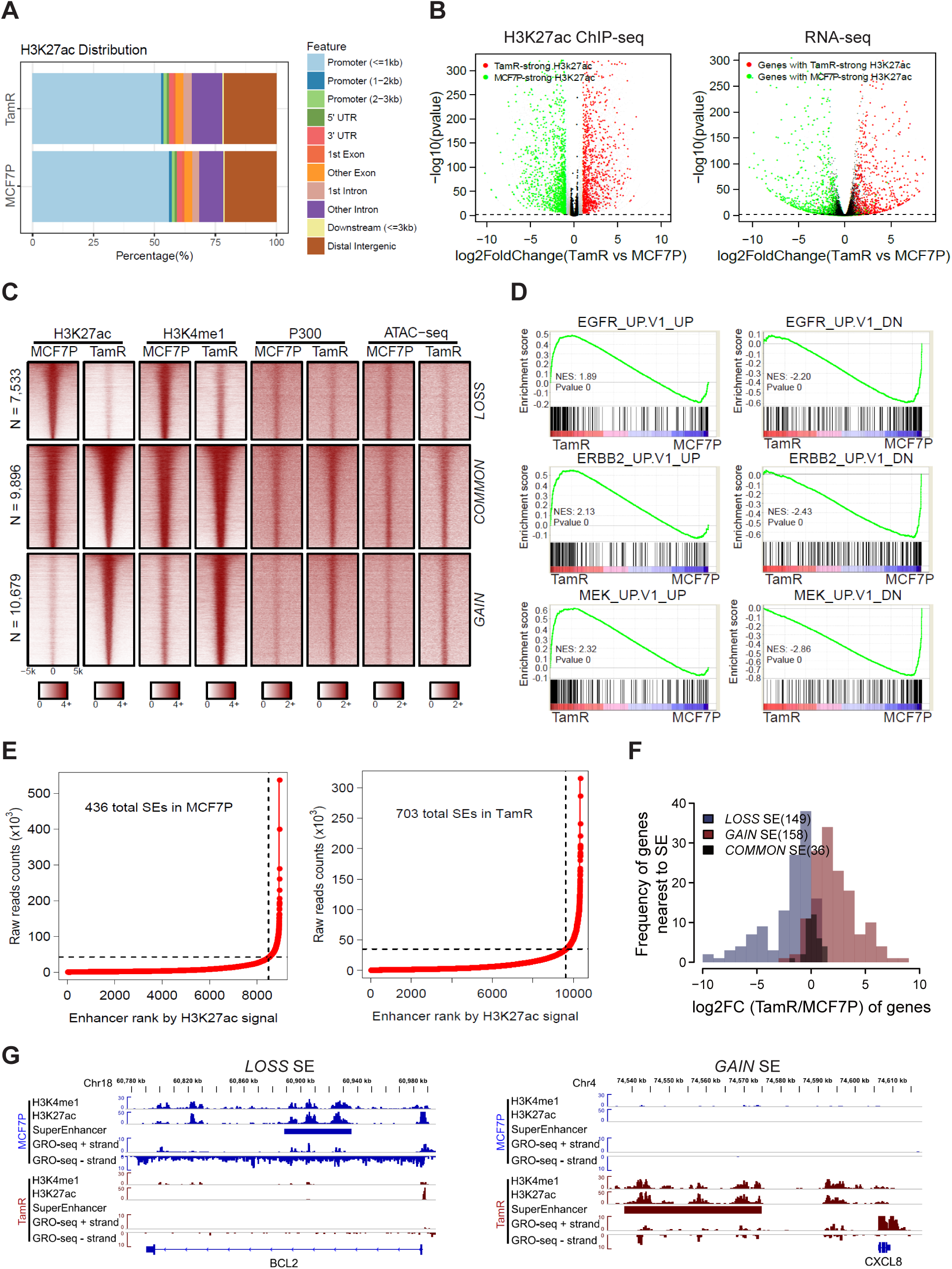
Endocrine resistance is associated with global enhancer reprogramming that drives plasticity-enhancing gene expression, Related to Figure 2. (A) Genomic annotations of the H3K27ac ChIP-seq signals in TamR and MCF7P cell lines. (B) Volcano plots showing the changes of H3K27ac signals at promoter regions correlate well with the changes in gene expression detected by RNA-seq in TamR cells. (C) Heatmap of H3K27ac, H3K4me1 and P300 ChIP-seq data for all identified lost, common and gained enhancers genome wide. Chromatin accessibility profiled by ATAC-seq at the corresponding genomic regions is also shown on the right. (D) GSEA analyses on RNA-seq data showing the enrichment of oncogenic signatures from MSigDB database in TamR or MCF7P cells. (E) Total super-enhancers (SEs) in MCF7P and TamR cell lines identified by the ROSE program ranked by H3K27ac signal intensities. (F) Histograms of the log2(Fold Change) of genes nearest to the differential SEs showing that gained SEs correlate with gene upregulation and lost SEs correlate with gene downregulation. (G) Genome browser snap images of lost SE at *BCL2* locus and gained SE at *CXCL8* locus. The SE gain/loss correlates well with gene upregulation and downregulation detected by GRO-seq.

**Figure S3.**
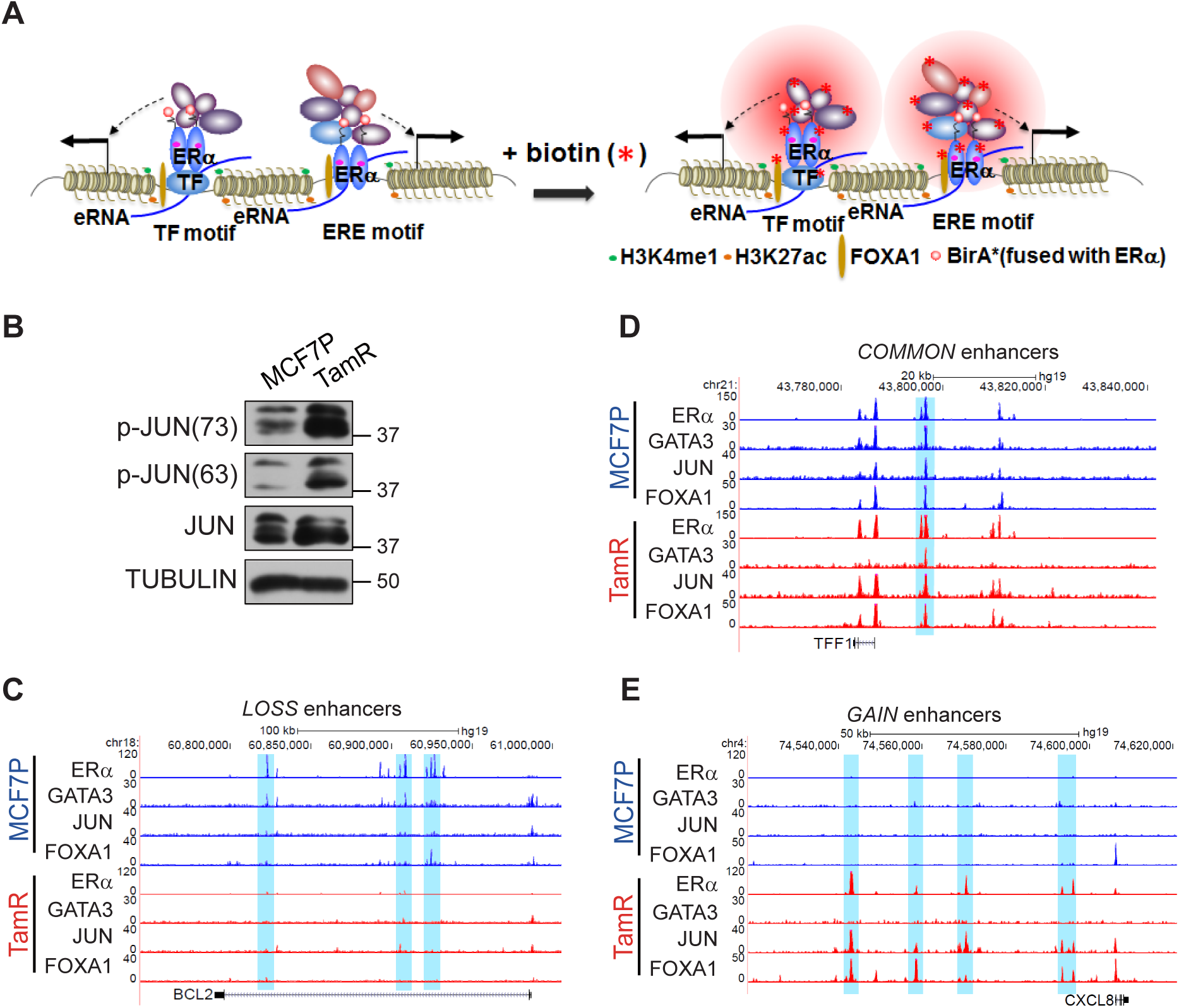
Altered interactions between ERα and GATA3/AP1 and their binding on ERα enhancers are associated with enhancer gain/loss reprogramming, Related to Figure 3. (A) Schematic diagram of BioID (*in vivo* proximity-dependent biotin identification) approach for identification of ERα-interacting nuclear proteins including both TFs and cofactors. (B) Western blot analyses of total JUN or phosphorylated JUN protein levels in MCF7P and TamR cells. Tubulin was used as a loading control. (C-E) Genome browser snap images of ChIP-seq data showing the co-binding of GATA3, JUN, FOXA1 and ERα at the *LOSS* enhancer regions near *BCL2* gene (C), the *COMMON* enhancer regions near *TFF1* gene (D), and *GAIN* enhancer regions near *CXCL8* gene (E) in both MCF7P and TamR cell lines.

**Figure S4.**
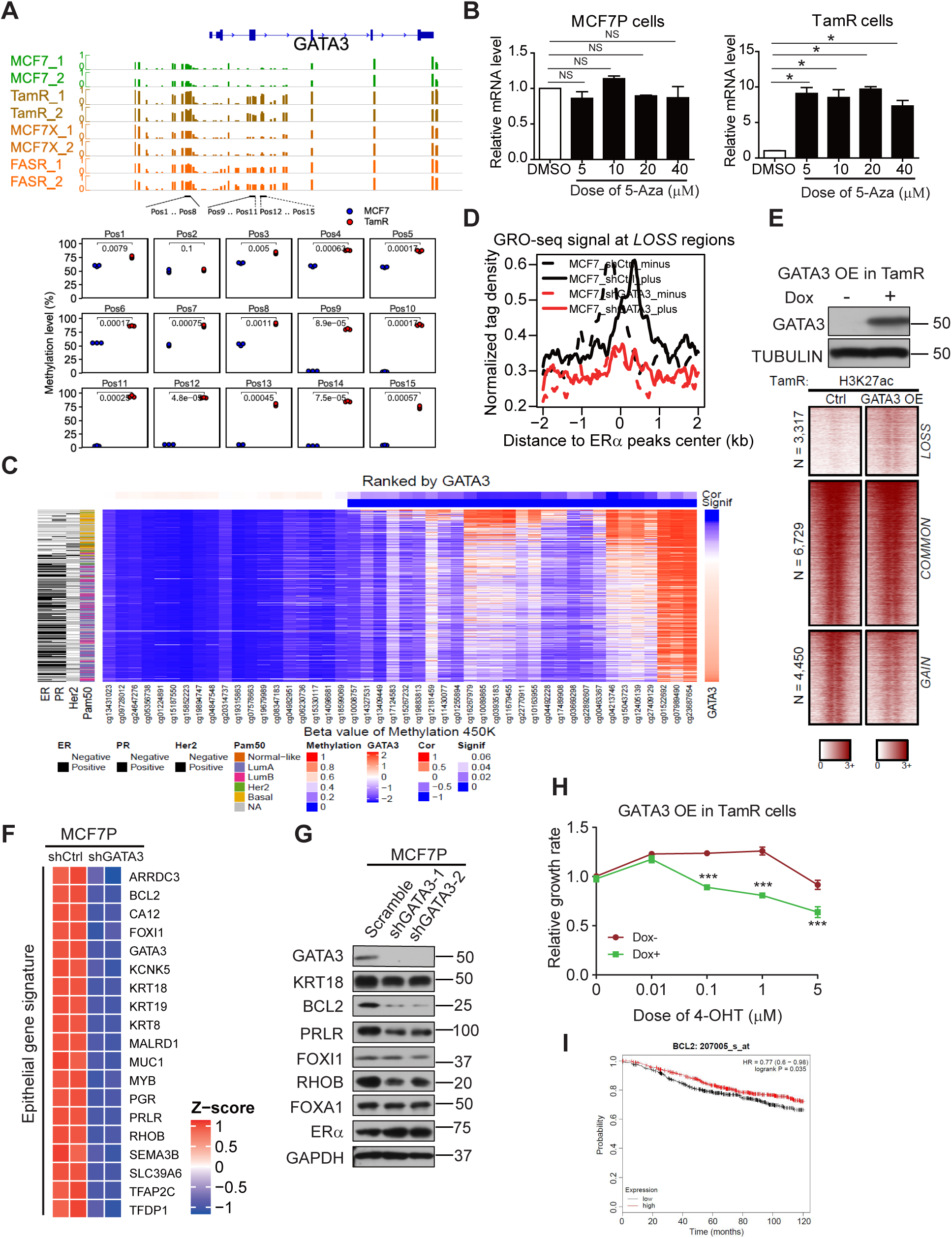
GATA3 is required for maintenance of *LOSS* enhancers and expression of epithelial makers, Related to Figure 4. (A) Our pyrosequencing analyses (bottom) showing a significantly higher level of DNA methylation at *GATA3* gene locus in TamR cells than in MCF7P parental cells used in this study. This is consistent with our analyses on *GATA3* locus (top) using published genome-wide DNA methylation data from three different pairs of endocrine-resistant MCF7-derived cell lines: tamoxifen-resistant (TamR), fulvestrant-resistant (FASR), and estrogen deprivation-resistant (MCF7X) cells ^39^. Thus, DNA-methylation mediated gene silencing at *GATA3* locus could be a general event during the development of endocrine resistance. (B) RT-qPCR showing transcript levels of *GATA3* in MCF7P and TamR cell lines with or without 5-Aza treatment for 100 hours. DNA demethylation induced by 5-Aza caused upregulation of GATA3 in TamR cells. Data were presented as means ± SEM. NS, not significant. *P <0.05 by 2-tailed t test. (C) Heatmap generated by integrating TCGA data on GATA3 mRNA expression level, DNA methylation, and breast cancer subtype. DNA methylation signals detected using specified DNA methylation probes (column) for every patient (row) negatively correlate with GATA3 expression levels. The heatmap matrix from top to bottom was sorted based on the GATA3 expression level from low to high. The results indicate that high DNA methylation and low GATA3 expression are associated with breast cancers that are negative to ER, PR, and HER2 and are often basal subtype. (D) Aggregate plots of the normalized GRO-seq tag density at *LOSS* enhancers in MCF7P cells transduced with shCtrl or shGATA3 lentiviruses. GATA3 knockdown greatly reduces eRNA transcription due to enhancer inactivation. The dashed line represents the minus strand and solid line indicates the plus strand of eRNA. (E) Heatmap of H3K27ac ChIP-seq data at *LOSS*, *COMMON* and *GAIN* enhancers (bottom) showing that overexpression of GATA3 in TamR cells can activate the silent *LOSS* enhancers. Western blot confirms the doxycycline-induced expression of GATA3 in TamR cells (top). Tubulin was used as a loading control. (F) Heatmap depiction of the downregulation of the epithelial differentiation related genes after knocking down GATA3 in MCF7P cells. (G) Western blot images of the indicated epithelial markers in MCF7P cells transduced with a scramble control or two different lentiviral shRNAs for GATA3 showing that GATA3 is required for the expression of these epithelial markers. (H) Overexpression of GATA3 in TamR cells resensitizes them to 4-OHT. CCK8 assays with a TamR stable cell line expressing doxycycline-induced GATA3 (-dox was used as control) were performed to check the relative cell viability of cells after treatment with indicated 4-OHT concentrations for 5 days. Data are presented as mean ± SEM. *** P < 0.001. (I) Relapse free survival (RFS) curves according to *BCL2* gene expression levels in patients receiving endocrine therapy. The curves were generated using data from kmplot website.

**Figure S5.**
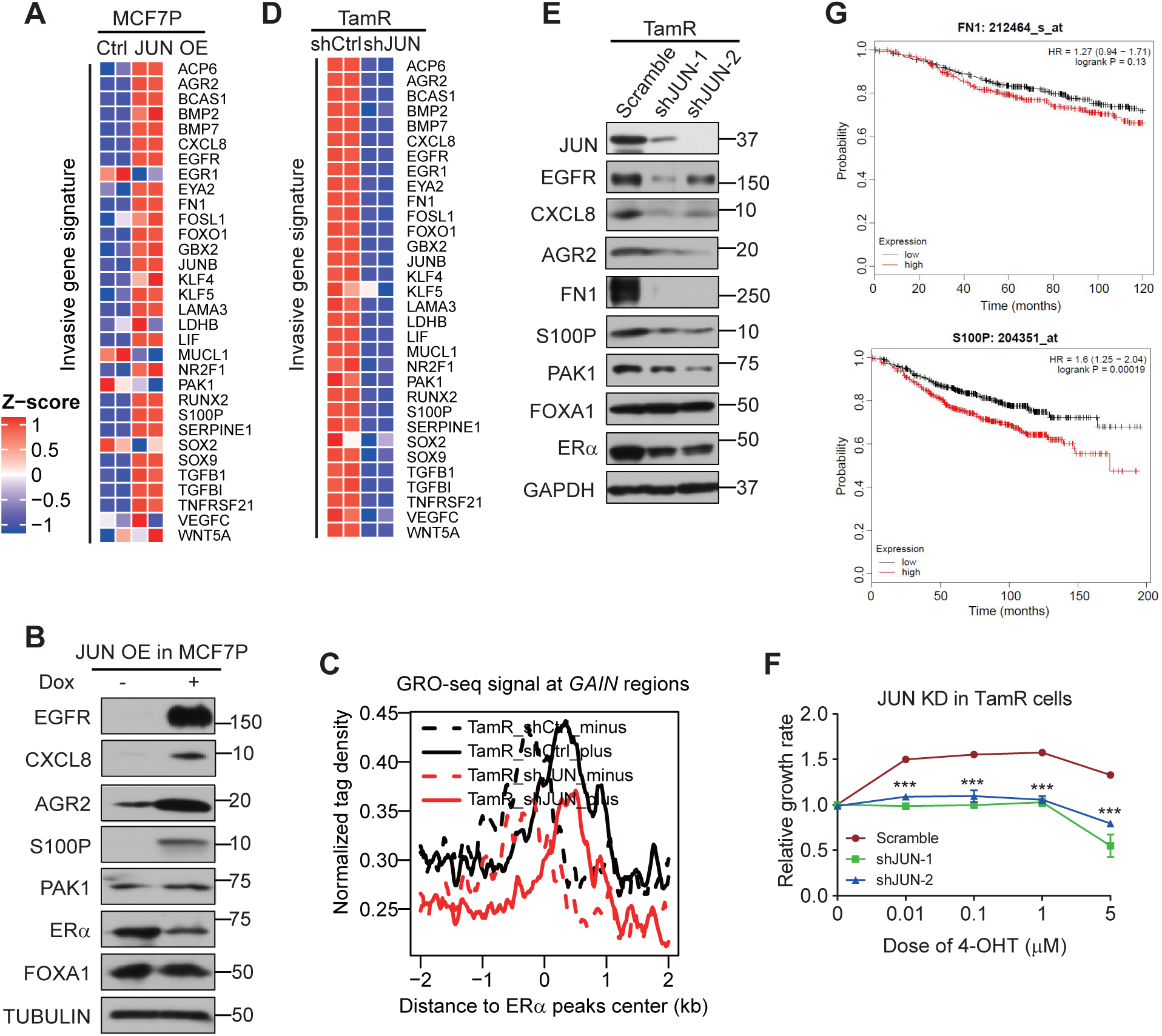
AP1-mediated *GAIN* enhancer activation promotes hormone resistance-associated gene program and phenotypes, Related to Figure 5. (A) Heatmap depiction of the upregulation of indicated invasive genes after JUN overexpression in MCF7P cells. (B) Western blot images of indicated invasive markers in MCF7P cells with or without JUN overexpression, showing that JUN overexpression is sufficient to activate the expression of these invasive markers. (C) Aggregate plots of the normalized GRO-seq tag density at *GAIN* enhancers in TamR cells transduced with shCtrl or shJUN lentiviruses showing that knockdown of JUN greatly reduces eRNA transcription due to enhancer inactivation. The dashed and solid lines represent the minus and plus strands of eRNA respectively. (D) Heatmap depiction of the downregulation of indicated invasive genes after JUN knockdown in TamR cells. (E) Western blot analyses on indicated invasive markers in TamR cells transduced with a scramble control or two different lentiviral shRNAs for JUN, showing that JUN is required for the expression of these invasive markers. (F) Knockdown of JUN in TamR cells resensitizes them to 4-OHT. TamR cells were stably knocked down with shJUN (a scramble shRNA was used as control) and CCK8 assays were used to check the relative cell viability of cells after treatment with indicated 4-OHT concentrations for 5 days. Data are presented as mean ± SEM. *** P < 0.001. (G) Relapse free survival (RFS) curves according to *FN1* and *S100P* gene expression levels in patients receiving endocrine therapy. The curves were generated using data from kmplot website.

**Figure S6.**
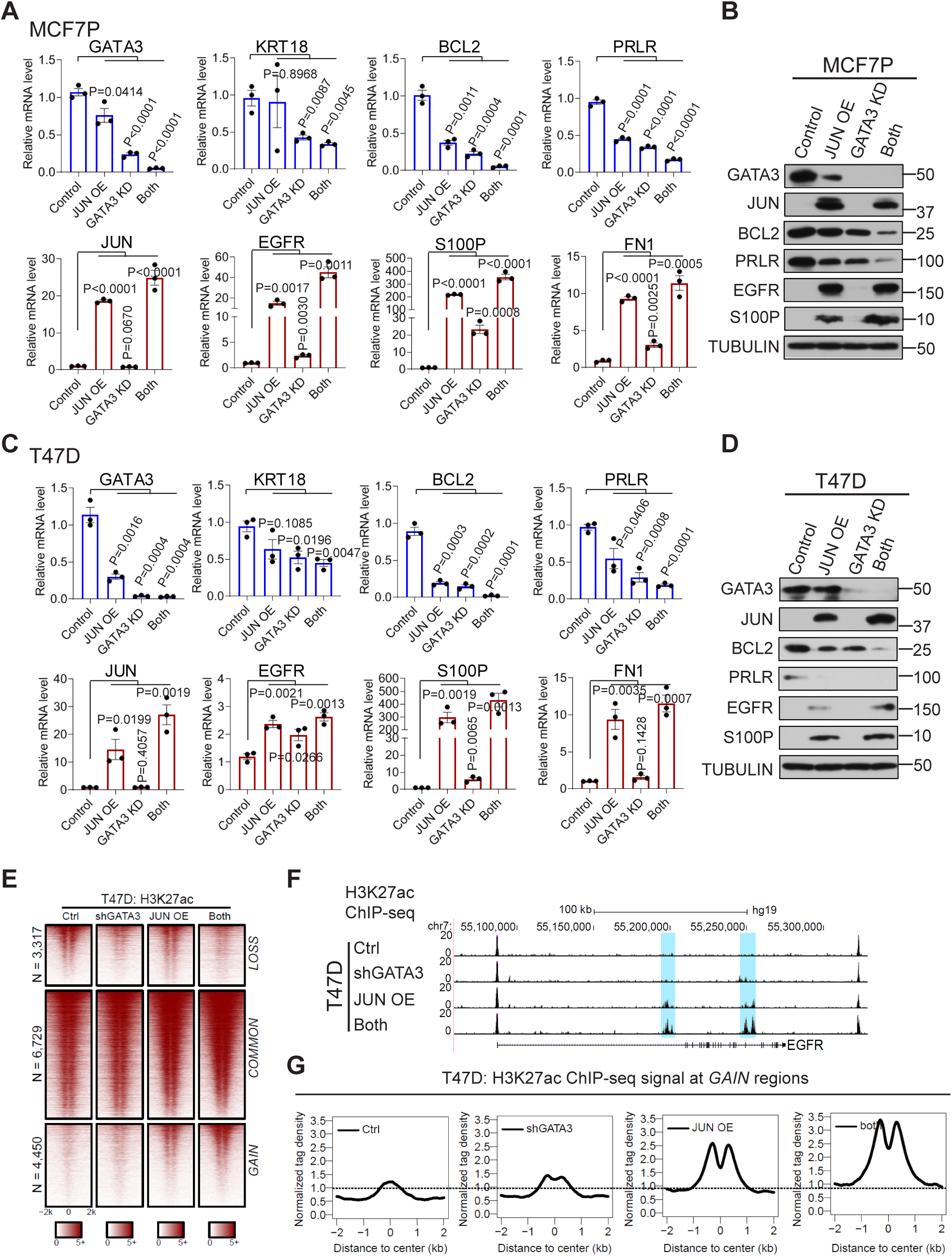
Coordinate role of GATA3 and AP1 in promoting gene expression alteration, Related to Figure 6. (A-B) RT-qPCR and western blot analyses of selected epithelial markers and invasion-related genes in MCF7P cells with indicated manipulations, showing the coordinate gene regulation effects by GATA3 and JUN. (C-D) RT-qPCR and western blot analyses of selected epithelial markers and invasion-related genes in indicated T47D cells with indicated treatments, showing the coordinate role of GATA3 and JUN in regulating gene expression. (E) Heatmaps of H3K27ac ChIP-seq data at *LOSS*, *COMMON* and *GAIN* enhancers in T47D cells with the indicated treatments. (F) Genome browser snapshot of H3K27ac ChIP-seq signals at the *EGFR* gene locus. GATA3 KD and JUN OE show a synergistic effect in T47D cells. (G) The aggregate plots of the normalized tag densities of H3K27ac ChIP-seq data at *GAIN* enhancers in T47D cells with indicated treatments. GATA3 KD and JUN OE demonstrate a synergistic effect on activating *GAIN* enhancers.

**Figure S7.**
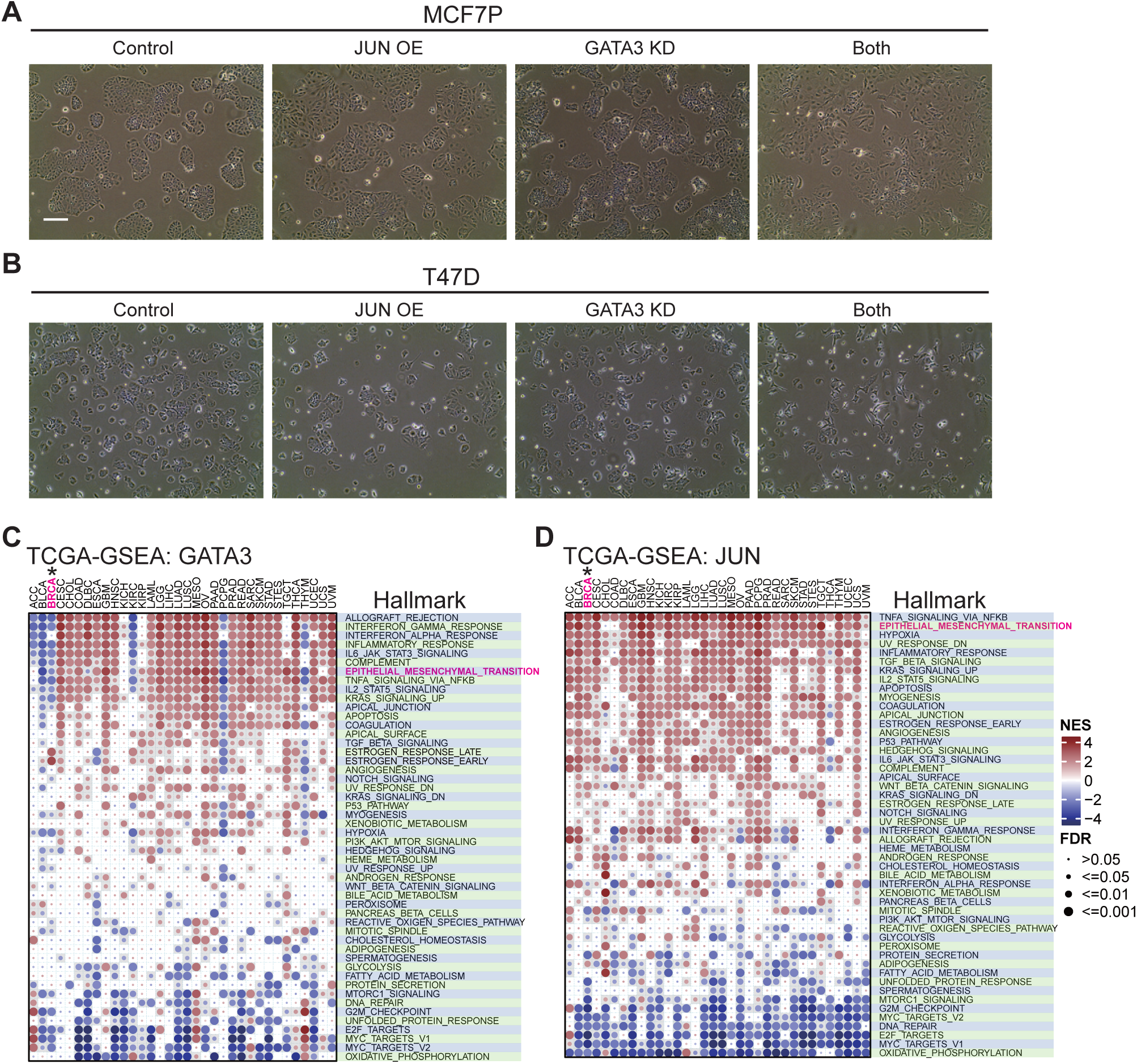
Combined effect of GATA3 and AP1 in enhancing breast cancer invasive progression, Related to Figure 7. (A-B) Representative brightfield pictures of MCF7P cells (A) and T47D cells (B) with indicated manipulations. The control cells display a typical epithelial cell-like morphology and grow in tightly packed clusters. Cells with both GATA3 knockdown and JUN overexpression have become more spread out (a phenotype of more invasive cancers) than the control and the cells with individual manipulation. Magnification, ×100. Scale bar, 100 µm. (C-D) GSEA analyses of RNA-seq data for 34 different cancer types including breast cancer (BRCA) from TCGA database showing the correlation of GATA3 and JUN expression levels with the enrichment of cancer hallmark gene sets from MSigDb database. We found that high expression level of JUN was positively associated with the enrichment of EMT pathway in breast cancer, however high expression level of GATA3 was negatively correlated with EMT pathway in breast cancer. The circle size indicates significance level; and the color represents the normalized enrichment score (NES).

